# “Distinct roles of dorsal and ventral subthalamic neurons in action selection and cancellation.”

**DOI:** 10.1101/2020.10.16.342980

**Authors:** Clayton P. Mosher, Adam N. Mamelak, Mahsa Malekmohammadi, Nader Pouratian, Ueli Rutishauser

## Abstract

The subthalamic nucleus (STN) supports action selection by inhibiting all motor programs except the desired one. Recent evidence suggests that STN can also cancel an already selected action when goals change, a key aspect of cognitive control. However, there is little neurophysiological evidence for a dissociation between selecting and cancelling actions in the human STN. We recorded single neurons in the STN of humans performing a stop-signal task. Movement-related neurons suppressed their activity during successful stopping whereas stop-signal neurons activated at low-latencies regardless of behavioral outcome. In contrast, STN and motor-cortical beta-bursting occurred only later in the stopping process. Task-related neuronal properties varied by recording location from dorsolateral movement to ventromedial stop-signal tuning. Therefore, action selection and cancellation coexist in STN but are anatomically segregated. These results show that human ventromedial STN neurons carry fast stop-related signals suitable for implementing cognitive control.

## Introduction

The ability to cancel plans that no longer support our goals is a critical feature of cognitive flexibility. In the motor system, this commonly manifests in the ability to cancel or inhibit actions already underway, such as stopping a movement towards a target. Computational models posit that brain circuits that inhibit actions (so called “stop pathways”) compete with circuits that promote actions (“go pathways”) and therefore an action can only be cancelled if activity in the stop pathway outraces activity in the go pathway (Logan & Cowan, 1984; Boucher et al., 2007; Schall et al., 2017; Schmidt et al., 2013). Physiologically, an instantiation of a go pathway in the human brain is the “direct pathway” that links motor cortex with the motor thalamus via the striatum and substantia nigra in the basal ganglia. Activation of this pathway ultimately transiently releases motor thalamus from its state of constant inhibition, which allows a movement to occur. A neurophysiological instantiation of a stop pathway that cancels movements when goals change has remained largely elusive, though is thought to involve connectivity between frontal cortex and some access point in the basal ganglia.

Recent evidence suggests that the subthalamic nucleus (STN) is the missing link (Jahanshahi et al., 2015; Aaron et al., 2014; Aaron et al., 2003; Aaron & Poldrack, 2006; Aaron et al., 2016; Obeso et al., 2014; Wiecki and Frank, 2013; Wessel et al., 2016a; Alegre et al., 2013). The STN excites the final output nuclei of the basal ganglia (globus pallidus internus, GPi; substantia nigra pars reticulata, SNr), thereby increasing the inhibition of the motor thalamus (Jahanshahi et al., 2015b). Classically, the STN is placed in the “indirect pathway” through the basal ganglia. This multisynaptic pathway is slow compared to the direct pathway and is thought to play a role in terminating actions that have already been completed by the go-pathway, or in inhibiting unwanted motor programs that compete with the go-pathway (Mink, 1996; Nambu et al., 2002). Recently, a monosynaptic “hyperdirect pathway” from frontal cortex to STN has been identified (Haynes & Haber, 2013; Chen et al., 2020), leading to the hypothesis that frontal cortex (specifically right inferior frontal gyrus) can bypass the slower multisynaptic inhibitory pathway to directly activate STN even more quickly than the go-pathway (Aron et al., 2003; Aron & Poldrack, 2006; Aron et al., 2014; Chen et al., 2020; Wessel et al., 2016a). If this hyperdirect pathway is activated, it would lead to a widespread increase in inhibitory drive and rapidly cancel an action when a goal quickly changes (Wessel and Aaron, 2017; Schmidt et al., 2013).

While the cytoarchitecture of the STN is largely homogenous (Yelnik and Percheron, 1979), different parts of STN have been segregated into distinct zones that contain different somatotopic maps and/or which receive input from different parts of cortex or basal ganglia (Keuken et al, 2012; Rodriguez-Oroz et al., 2001; Nambu et al., 1996; Iwamuro et al., 2017; de Long et al., 1985). Anatomical tracing studies in nonhuman primates reveal that the hyperdirect stop-pathway from frontal cortex terminates in the ventral STN, while motor cortex projects to dorsal STN. These observations imply that STN is divided into different functional zones (Haynes & Haber, 2013). This leads to the overall hypothesis that dorsal regions support action selection and motor control, whereas ventral regions support higher cognitive functions such as stopping (Greenhouse et al., 2011; Jahanshai et al. 2015; Mallet et al., 2007; Eweret et al., 2018). While electrical stimulation and local field recordings support this functional parcellation of action selection and cancellation (Chen et al., 2020), there is at present no single neuron evidence for this hypothesis in humans.

If the STN has distinct functional zones that play different roles in movement and cognition, this would be of profound clinical significance. The STN is a routine target for deep brain stimulation (DBS) that alleviates the symptoms of movement disorders such as Parkinson Disease (Pouratian et al., 2012). A common and undesirable side-effect of such stimulation is inappropriate impulsive behavior, which empirically has been associated with the placement of electrodes in ventral STN (Wouwe et al, 2017; Chen et al., 2020; Jahanshahi et al., 2015ab; 2015; Greenhouse et al., 2011; Hershey et al, 2010). It has been hypothesized that high frequency electrical stimulation at these more ventral contacts impinges on the ability to cancel actions and thereby leads to impulsive behaviors. Electrical stimulation at more dorsal sites may lead to better improvement in motor control with fewer cognitive side effects (Greenhouse et al., 2011). Additionally, if neurons that support stopping have distinct firing features from movement neurons, this information could be utilized to design stimulation protocols that specifically inhibit specific cell types which would be a major clinical advance.

It remains unknown whether this functionally-segregated model of STN, which is contrary to the standard model of STN as monolithic relay station that plays no role in cognition, is applicable to the human brain. Here, we recorded individual neurons in STN as humans performed a stop-signal task to determine (1) are there neural representations of stop or go pathways in the STN and (2) do these representations localize to distinct subfields of STN as would be predicted by a frontal-cortical hyperdirect pathway.

## Results

### Task and Behavior

We recorded single neurons from the human STN and field potentials (ECoG) from sensorimotor cortex as subjects performed a stop signal task during surgery for implantation of DBS electrodes to treat symptoms of Parkinson Disease (33 surgeries/sessions in 19 subjects, see Methods, Figure 1, Table S1). Subjects were instructed to move a joystick as quickly as possible in the direction of a go-signal (white arrows, Figure 1AB) using the hand contralateral to the recording site. On 1/3 of trials, subjects were presented with an instruction to stop (or cancel) the planned action. The onset of the stop signal occurred at a variable time delay relative to stimulus onset. It was set at the beginning of the session to 300 ms and was adjusted in ±/-50 ms increments so that subjects successfully stopped on ^~^ 50% of trials.

**Figure 1:**
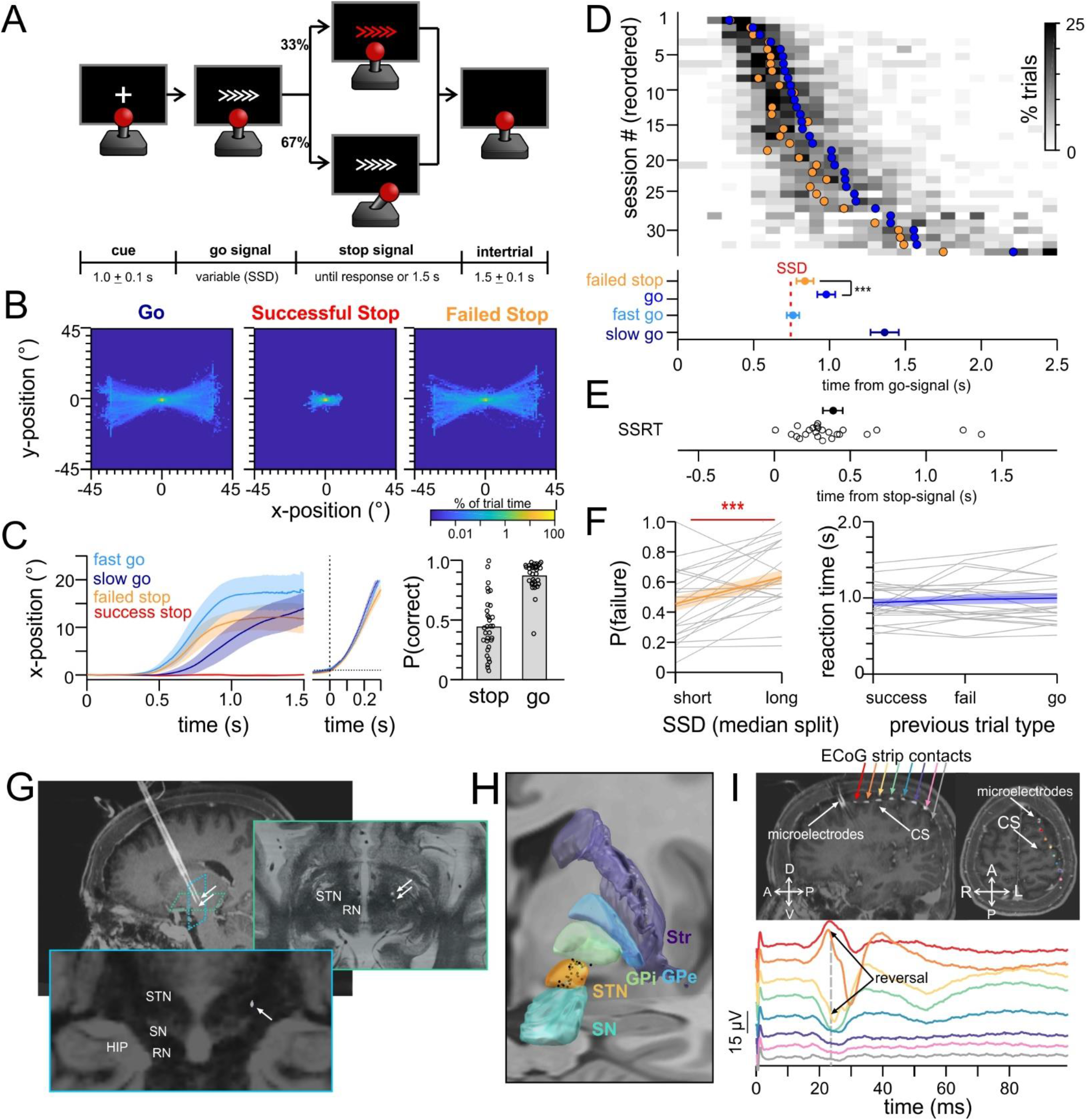
Stop-signal task and recording setup. **A.** Task. On each trial a go-signal (white arrows) indicated the direction the subject should move the joystick. On a random subset of trials, a stop-signal (red arrows) appeared at a variable delay (stop signal delay, SSD). If the stop-signal appeared, the goal of the subject was to remain still, ignoring the prior go-signal. **B.** Heatmap of joystick position across all subjects (negative x-position = subject moves left). **C.** Time-course of joystick trajectory aligned to go-signal onset (left) and movement onset (middle inset). Horizontal dotted line indicates threshold for movement detection. Bar plot shows probability of successfully stopping on a stop trial and probability of moving in the correct direction on a go trial (each dot is a session). **D.** Distribution of reaction times. Each row represents the behavior during a single session. Gray heatmap shows binned reaction times across all trials (histogram). Filled circles represent the mean response times on failed stop trials (orange) and on successful go trials (blue). Population averages and s.e.m. are depicted at bottom for each trial type. **E**. Scatterplot at bottom shows the distribution of the stop-signal reaction times (SSRT). **F.** (left) Inhibition functions show the probability of failing to stop on trials with short or long SSDs (median split). (right) Absence of sequence effects. The reaction time (RT) on go-trials that follow a successful stop, a failed stop, or another go-trial. Gray lines are individual subjects. Blue and orange lines depict mean ± s.e.m. **(F-H**) Recording setup. **F.** Intraoperative CT scan fused with pre-operative MRI showing two microelectrodes targeting STN. Arrows indicate microelectrode tip in STN. **G.** Reconstruction of STN microelectrode recording sites, displayed in dorsal-ventral and medial-lateral space with respect to patient midline and anterior-posterior commissure. Each black dot is the location of a recorded neuron. **H.** Intraoperative CT scan fused with pre-operative MRI showing the location of the ECoG strip placed over sensorimotor cortex. Traces show the sensorimotor evoked potential elicited by median nerve stimulation. Each color indicates a different ECog contact. *p<0.05, **p<0.01, ***p<0.001

On average, subjects moved the joystick in the correct direction on 87±11% of go trials and successfully stopped on 44±25% of stop trials (Figure 1C). In some sessions (n=9), subjects performed fewer than 5 successful or failed stop trials; these sessions were excluded from stop-related analysis (mean stopping accuracy: 14±12% for subjects with <5 successful stops, 90±13% for subjects with <5 failed stops). In the remaining sessions, subjects successfully stopped on 45±19% of stop trials and chose the correct direction on 91±7% of go trials. Throughout the paper we analyze four types of conditions: “fast-go trials” (trials where a go-signal was followed by a fast reaction time and the subject moved in the correct direction), “slow-go trials”, “successful-stop trials” (trials where a stop signal appears and the subject successfully cancels their action) and “failed-stop trials.”

Under a race model of stopping, trials with short-latency activation of motor plans are more difficult to cancel than trials in which the motor plan is initiated more slowly. This model predicts that reaction times on failed-stop trials should be skewed toward the faster tail of the go reaction time distribution. Confirming this prediction, we found that reaction times on failed-stop trials were shorter than the average reaction time on go-trials (mean difference in reaction time=140±10 ms; paired t-test t(23)=-7.01 p=3.8 x 10^−7^) (Figure 1D). The two key parameters in race models are the SSD (“stop signal delay”, time between stimulus onset and onset of stop signal) and the SSRT (“stop signal reaction time”, the latency of the stop process). The average SSD across the sessions was 480±236 ms. In each session we split the variable stop signal delays (SSDs) into short and long durations (median split) to estimate the inhibition function of each session (Figure 1F, left panel). Longer SSDs significantly increased the probability of failing to stop, in line with a race model of action cancellation (comparison of short vs long SSD: paired t-test t(22)=3.75, p=0.0011; average P(respond) for short SSD=0.46±0.23; average P(respond) for long SSD=0.63±0.22). Based on the distribution of reaction times and the rate of successful stopping, we estimated the stop-signal reaction time (SSRT) to be 361±334 ms (Figure 1E, integration procedure, see Methods; Verbruggen et al., 2019). Lastly, we tested whether reaction times depended on the behavior in the previous trial, to see if subjects delayed their responses after a failed stop trial. We found no such dependence: on average, subjects responded with similar reaction times on go-trials that followed successful or failed stop trials (Figure 1F, right panel, paired t-test for reaction times following successful vs. failed stop: t(23)=0.88, p=0.388; successful stop vs. go: t(23)=1.09, p=0.285; failed stop vs. go: t(23)=0.56, p=0.579).

### Neurons in dorsal STN were activated prior to and during the execution of movements

We recorded a total of n=83 well isolated and stable neurons in STN while subjects performed the stop-signal task (Figure 1GH show recording sites, on average 2.5±2.4 cells were recorded per session; Supplementary Figure 1 shows sorting quality metrics). The responses of n=32/83 neurons (39%) were significantly modulated when subjects moved the joystick (see example units in Figure 2A) (39% significantly greater than chance=8±3%; 500 shuffled data trials, t(499)=31.8, p=4.4 x 10^−122^). These “movement-related” neurons had an average baseline firing rate of 13.2±17.3 Hz, responded robustly around movement onset (average peak firing rate: 18.3±19.4 Hz, 58% increase in firing rate, mean effect size=0.37±0.29 s.d.) and the majority (30/32, 94%) signaled movement onset with an increase in firing rate. Subjects responded with variable latencies following the go-signal, allowing us to align the neural activity to either the onset of the go-signal or the onset of the movement to assess which event better predicted neural activity. On average the neural activity was more tightly time-locked to the onset of the movement, suggesting that these neurons play a role in movement selection, execution, or sensorimotor feedback rather than evaluating the go-signal itself (Figure 2B, paired t-test comparison of peak firing rate aligned to movement vs. target, t(31)=3.980, p=0.000387).

**Figure 2:**
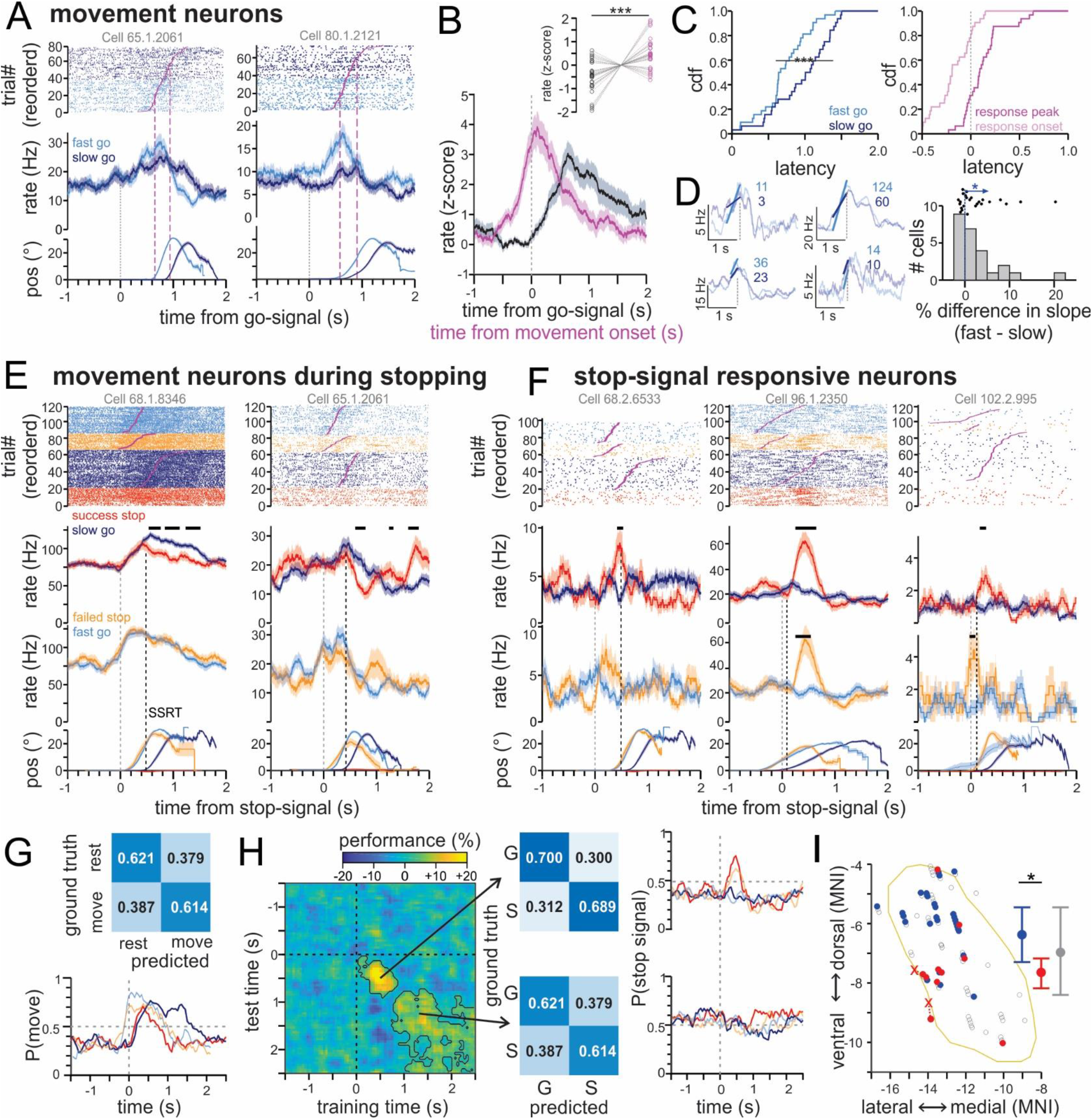
Single neuron responses during the stop signal task. **A.** Raster-plots and peri-stimulus time histograms (PSTHs) of two example STN neurons that increased their firing rates during movement. t=0 is onset of the go-signal. Trials are grouped by fast (light blue) and slow (dark blue) response times. Bottom trace shows average joystick position along the x-axis. **B.** Average normalized firing rate of all movement neurons, aligned to onset of the movement (pink) or the go-signal (black). Inset shows peak firing rate attained for each neuron for the two temporal alignments. **C.** Response latency. (left) Time at which neurons attained their peak firing rate following the go-signal on trials with fast (light blue) vs. slow (dark blue) reaction time. (right) Time relative to movement onset that movement-related neurons began to respond (light pink) and attained peak firing rate (dark pink). **D.** Slope of firing rate increase during movement. (left) Average firing rate of four example movement-related neurons aligned to movement onset during fast (light blue) and slow go trials (dark blue). The fit line was used to estimate the speed of increase firing rate indicated in text (slope; units = Hz/sec). Light traces show PSTH, dark lines show fitted slope. Vertical dotted line=movement onset. (right) Difference in slope on fast vs. slow go trials for all movement-related neurons that began to fire before movement onset (n=25/32 movement neurons). **E.** Example activity of two movement-related neurons in response to the stop-signal. Raster plots, PSTH, and joystick position aligned to stop-signal onset for successful (red) and failed (orange) stop trials. Fast go (light blue) and slow go (dark blue) trials are aligned to the timepoint when the stop-signal would have appeared had the trial been designated a stop-trial. **F.** Example stop-signal responsive neurons. **(G-H**) Decoding results. **G.** Confusion matrix for classifier trained to decode movement from neural firing rates. Trace shows the probability the decoder predicted movement throughout the trial for each trial type. **H.** Performance of a classifier trained to discriminate stop-signal trials from go-trials and tested at different time points. t=0 is the time the stop-signal appeared (or for go-trials, the time it would have appeared). Contours outline periods where performance is significantly better than chance (shuffled data, cluster correction). Confusion matrices at two example time periods. Traces show the probability the decoder predicts a stop-signal at different time points during the trial. **I.** Anatomical location of movement (blue)and stop signal neurons (red) in STN (yellow outline). Recording sites were reconstructed using a post-operative CT scan. For illustration, we obtained an intraoperative image of the two micro-electrodes where stop-signals were recorded in one patient (Figure 1E). For this example, the red “X”’s show the location of the recordings inferred from the intraoperative image, connected with a dotted line to the location predicted by the post-surgical reconstruction. The more dorsally located “X” (near −8 dorsal-ventral MNI) is the location of the neuron shown in Figure 2F, far right panel. Error bars show mean ± 1^st^ and 3^rd^ quartile location of neurons along the dorsal-ventral axis. (AP slice=-14.06 mm) *p<0.05, **p<0.01, ***p<0.001

On average, movement-related neurons responded similarly to movements to the left or right (paired t-test comparing mean firing rate on left vs right movements for population of movement neurons, t(31)=0.696, p=0.492). We also compared firing rates during abduction (movement away from the body, move left with left hand, right with right hand) and adduction (toward the body). Adduction and abduction elicited similar changes in firing rates (t(31)=0.765, p=0.450). Thus, the responses of movement-neurons were not sensitive to movement direction.

The majority of movement-related neurons (25/32, 78%) began to fire before the earliest detectable movement of the joystick, suggesting a role in the early aspects of movement production, e.g., selection, planning, execution, as opposed to sensorimotor feedback. If movement neurons play a role in producing movements their responses should differ on trials with fast vs. slow reaction times. Neither the firing rate at baseline (500 ms prior to target onset), the firing rate at movement onset, nor the peak firing rate during movement was predictive of reaction time (t-test comparing mean firing rates for fast vs. slow go trials; baseline: t(31)=0.238, p=0.814; movement onset t(31)=0.862, p=0.395; peak rate t(31)=0.738, p=0.466). However, the time-course of activation revealed that when reaction times were faster, movement-related neurons attained their peak firing rates earlier in the trial (paired t-test comparing response latencies on fast vs slow trials: t(31)=3.876, p=0.000523, mean difference in neural response time=364±250ms) (Figure 2C, left panel). Computational models of movement execution suggest that movement-related neurons ramp to a threshold required for movement activation; when neurons ramp to this threshold more quickly the movement is executed earlier (Hanes and Schall, 1996). To examine such ramp-to-threshold activity, we fitted a line to the peri-stimulus time histogram of each neuron and calculated the slope between the timepoint when the neuron became active and when the movement occurred (least-squares regression). Traces from four example neurons show that the slopes were steeper on trials with fast compared to slow reaction times (Figure 2D, example traces). On average, the population of movement-related neurons had significantly steeper slopes on fast vs. slow reaction time trials (t-test comparing the slope on fast and slow trials for movement-neurons that had a response onset that preceded movement, t(24)=2.458, p=0.022) (Figure 2D). This time-course of activity suggests that the movement-related neurons played a role in selecting or executing movements, a feature of the go pathway. Other movement neurons whose activity increased *after* movement onset (n=7/32 neurons) may have played a role in monitoring proprioceptive feedback during the movement or updating the motor plan as the joystick was moved along its trajectory.

### Stop-signals activated neurons in ventral STN and reduced the activity of movement-related neurons

Of the 83 STN neurons we recorded, 47 neurons met the criteria required for subsequent analysis related to stopping (at least 5 successful-stop and 5 failed-stop trials). We first examined how the presentation of the stop signal modulated the activity of the movement-related neurons described above.

23/47 neurons were movement responsive (49%), as defined above. The activity of two example movement neurons in response to stop signals is shown in Figure 2E. These neurons had an elevated firing rate prior to movement onset (indicated by joystick trace at bottom) but this firing rate was reduced on successful stop trials compared to latency matched slow go-trials (compare red and dark blue lines Figure 2E). On failed stop trials, these movement-neurons were activated in a similar manner as latency-matched fast go trials. On average, movement neurons had lower firing rates on successful stop trials (paired t-test comparing mean firing rate on successful stop vs. slow go trials: t(23)=2.34, p=0.028). In contrast, their firing rate did not differ significantly between failed stop trials and fast go trials (paired t-test comparing mean firing rate on failed stop vs. fast go trials: t(23)=1.39, p=0.177). This is consistent with a role in movement: when a movement was cancelled after a stop signal, these neurons reduced their activity (comparison of activity on successful stop vs. slow go trials). If a subject moved after a stop signal (i.e., was unsuccessful at cancelling the action), the activity was similar to that of latency matched fast go trials.

To assess the response dynamics across the population of 47 neurons, we trained a classifier to differentiate between periods of rest and movement using the firing rates of the population of neurons during a single trial (see methods, single trial population decoder) (Figure 2G). The decoder differentiated periods of movement from rest (500 ms pre-target baseline) with 66 % accuracy, significantly better than chance (chance performance based on shuffled bootstrapped labels= 50±1%; t-test comparing decoder accuracy to bootstrapped data: t(999)=1485, p=1.8 x 10^−1672^). While the decoder had no a priori knowledge about the trial type (stop or go) or trial outcome (success or failed stop), the decoder nevertheless predicted movement to occur earlier on fast go trials compared to slow go trials (compare P(movement) for light vs dark blue traces) (Figure 2G). Also, on successful stop trials the probability of decoding movement decreased following the stop signal but was elevated for all other trial types (red trace, Figure 2G). Thus, the population activity of STN differentiated trials when movements were executed from those when movements were cancelled.

We next examined whether any neurons responded to the stop-signal. Of the 47 neurons, 10 (22%) exhibited significant changes in firing rate following the onset of the stop signal compared to a pre-stop signal baseline (22% significantly greater than chance=5±3%; 500 shuffled data trials, t(499)=142.3, p=4.4 x10^−406^). The onset of these “stop-signal responsive neurons” was 391±279 ms, within the time-frame of the SSRT (mean SSRT= 361±334 ms; two sample t-test comparing stop-signal response latencies and SSRTs: t(33)=0.253, p=0.802). Figure 2F shows three example neurons that increased their firing rate to the stop signal. We next compared the responses of neurons on stop trials compared to latency matched go trials. Five neurons (11%) significantly differentiated successful stop trials from slow go trials and failed stop trials from fast go trials (4/5 of these neurons also met classification for stop-signal responsive; 11% significantly greater than chance=1±1%; 500 shuffled data trials, t(499)=9.5, p=1.9 x 10^−19^). These neurons represent a subset of stop-signal neurons that registered a stop-signal regardless of trial outcome (success or failure to stop) and are compatible with those expected from neurons in the stop pathway. Two stop-signal responsive neurons (4%) only differentiated successful stop trials and one (2%) only differentiated failed stop trials (see examples of other responses in Supplementary Figure 2). Some of the stop-responsive neurons (n=7) also met criteria to be classified as movement responsive, suggesting some overlap between stop and go-pathways within the STN.

To better understand how the population of neurons responded to stop signals, we trained a single-trial classifier to differentiate between trials with and without a stop signal (similar to how we trained a decoder to decode when movement occurred, see above). The heatmap in Figure 2H shows the performance of the decoder with contour lines outlining areas where the decoder performed significantly better than chance (chance determined by training decoders with shuffled trial labels see Methods). The decoder differentiated stop trials from go trials during an early (less than 500 ms) and late period (> 1 s) after the appearance of the stop-signal trial. To better understand the decoder performance, we plotted the prediction of the decoder for each trial type using these two different training periods. Shortly after the stop-signal the probability of decoding a stop signal rapidly increased, regardless of the outcome (success or fail) of the stop trial (Figure 2H, top traces). This result suggests that the STN registered stop-signals, irrespective of the eventual motor output, which is a signal required to initiate cancelling actions. In contrast, the late component (Figure 2H, bottom traces) primarily differentiated successful stop trials from other trial types, indicating that this late component reflects regulation of the successful termination of the stopping process.

Lastly, we compared the anatomical position of movement and stop-signal neurons to test the hypothesis that different parts of STN participate in different behaviors. Stop-signal neurons in the STN were located significantly more ventral in the STN than movement-related neurons (Figure 2I; dorsal-ventral MNI coordinate of stop signal neurons= −7.6±1.5 mm; movement-related neurons= −6.4±1.4 mm; two-sample t-test t(41)=2.48, p=0.017). These data support the hypothesis that processes of action selection (movement) and action cancellation (stop-signal) are supported by neural populations located in distinct parts of the STN.

### Beta burst increases are too late to initiate stopping

Simultaneously with the single neuron recordings in STN we also recorded local field potentials (LFPs) from the microelectrode tip and iEEG signals from sensorimotor cortex (electrode localization using somatosensory-evoked potentials, see Methods, Figure 1I). As expected, field potentials from both STN and sensorimotor cortex exhibited prominent beta activity that was, on average, suppressed during the preparation and execution of a movement (Supplementary Figure 3) (Torecillos et al, 2018; Wessel, 2020; Feingold et al., 2015; Cagnan et al., 2019). However, beta activity in STN and motor cortex was not a continuous oscillation but rather occurred in bursts. We therefore next identified beta bursts (Torricellos et al., 2018) to investigate the time course of beta activity during the task and its relationship to single neuron spiking.

Beta bursts in motor cortex were 168±22 ms in duration and occurred 52.4±6.8 times per minute. In STN, bursts were 151±16 ms in duration and occurred 49.5±6.2 times per minute. As expected, beta bursting was pronounced in both motor cortex and in STN during the inter-trial interval and was reduced following onset of the target signal (Figure 3A). Similar to how cortical and STN beta power is often increased on trials where a subject successfully stops (see Supplementary Figure 3, Alegre et al., 2013; Benis et al., 2014; Wessel et al., 2016a; Fischer et al., 2017), beta burst probability was also higher on successful stop trials compared to latency matched go trials (Figure 3A, right panels).

**Figure 3:**
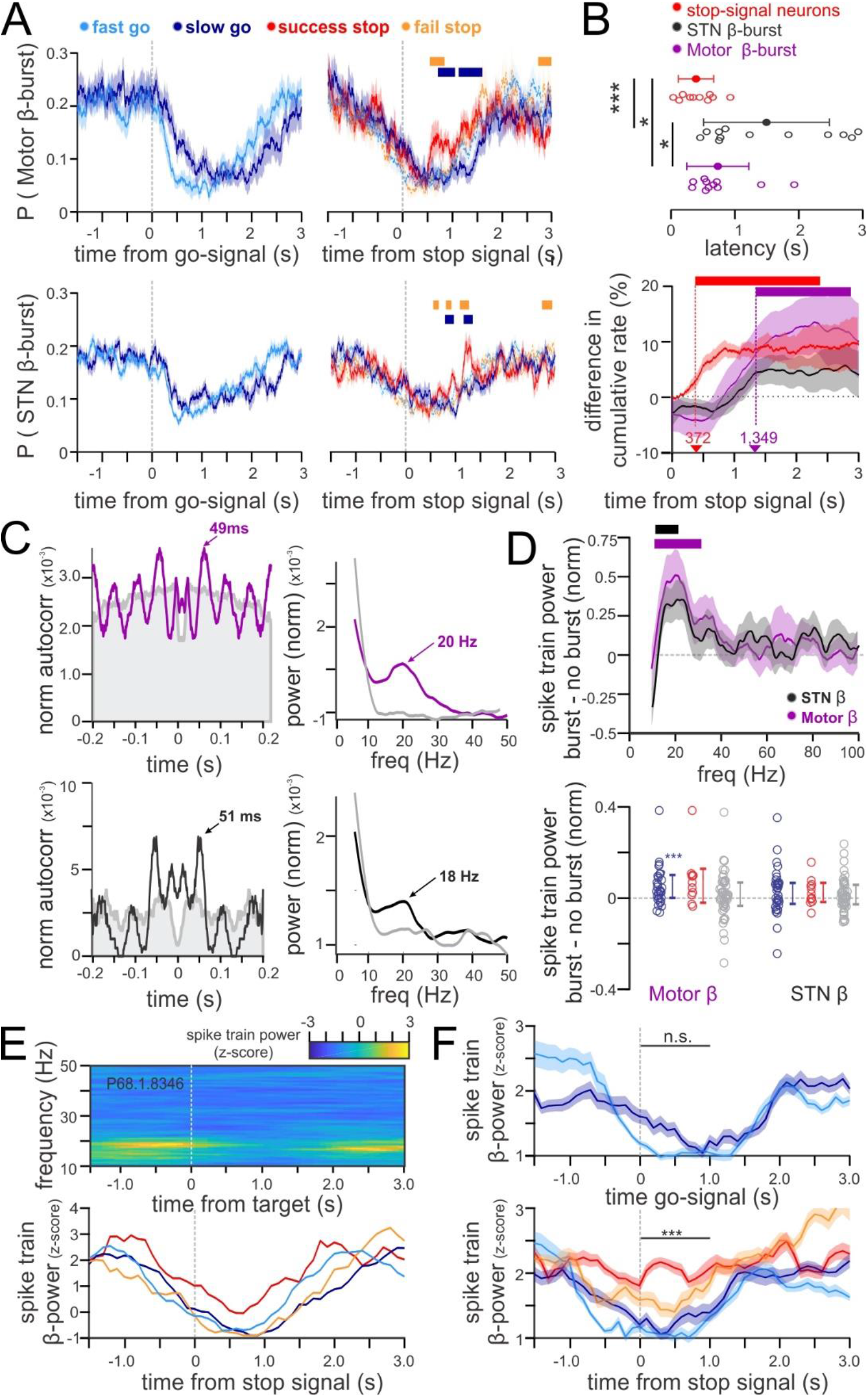
Beta activity of STN neurons changes after SSRT. **A.** Probability of a beta burst occurring in motor cortex (top) or STN (bottom) during the task. Left shows activity on go-trials aligned to the go-signal. Right shows activity aligned to the stop-signal. Horizontal bars indicate timepoints when beta burst probability significantly differed between successful stop trials and other trial types. Shading=s.e.m. **B.** Latency comparison between beta-burst onset and stop-related neuronal firing. (top) Estimated onset of stop-signal neurons and beta activity during successful stop trials. (bottom) Differential latency analysis comparing the response time of stop-signal neurons and beta bursting on successful stop vs. latency matched slow go trials. Shading=s.e.m. **C.** Autocorrelograms and power spectrums of the spike trains of two example STN neurons. The top neuron fired at a beta rhythm during beta bursts in motor cortex (purple) but not outside beta bursts (gray). The bottom neuron fired at a beta rhythm during STN beta bursts (black). **D.** (top) Average power spectrum of all neurons (n=83) during beta bursts in motor cortex (purple) and STN (black). Plots show the difference in power during vs. outside beta bursts (value=0 indicates beta burst does not predict beta spike firing; bars denote significance from 0; p<0.05) (bottom) The mean spike-train beta power for different types of neurons (gray=non-responsive, blue=movement-related, red=stop-signal). Power is calculated inside vs. outside beta bursts for motor cortex (left) and STN (right) **(E-F**) Increased beta-related firing during successful stop trials. **E.** Example neuron. (top) Power of the spike train at different time points following the go-signal (2000 ms bins, 100 ms sliding window). (bottom) Spike-train power in the beta band as a function of time for different trial types. **F**. Population summary. Average spike-train beta power across all 10 neurons that exhibited significant beta oscillatory activity. (top) activity on go-trials aligned to go-signal (bottom) activity on all trials, aligned to stop-signal. Shading=s.e.m. (significant p-value signifies difference between successful stop and latency matched go trials) *p<0.05, **p<0.01, ***p<0.001

However, the time at which beta burst probability increased relative to the stop-signal was relatively late (^~^ 1 second) calling into question whether this increased activity played an active role in stopping. To test this hypothesis, we calculated the earliest detectable difference in beta probability between successful stop and latency matched slow go trials for each session (cluster-wise statistic, identifying first bin with significant difference, p<0.05). STN beta burst probability increased 1,407±489 ms after the stop signal while ECog beta bursts increased 732±911 ms after the stop signal. This temporal latency was significantly later than the onset of stop-signal responses at the single-neuron level and significantly after the mean SSRT (Figure 3B) (two-sample t-test comparing latencies of stop neurons vs. STN beta: t(20)=3.39, p=0.0015; latencies of stop neurons vs. ecog beta: t(21)=2.18, p=0.020; latencies of STN beta vs SSRT: t(35)=5.13, p=0.000011; latencies of ECog beta vs SSRT: t(34)=2.659, p=0.012). Note that the onset of increased beta probability was not always quantifiable for every session and the method we used above used binned firing rates (see methods). As a control, we therefore also confirmed above findings using a different method: bin-free differential latency analysis (see methods; Xiang & Brown, 1998; Rutishauser et al., 2015). This method revealed that stop-signal neurons began to fire 372 ms after the stop-signal, whereas the probability of seeing an increase in motor cortex beta burst activity changed significantly at 1,349 ms. (Figure 3B; the differential latency for STN beta probability did not achieve significance so a latency could not be calculated). It is generally thought that an active stopping process needs to be initiated before the SSRT. Since our results indicate that beta burst increased occur significantly later than the SSRT, we conclude that modulations in beta activity were unlikely to be the mechanism that initiated stopping. Instead, we hypothesize that the stopping process was initiated by the stop-signal neurons described above.

We next turned our attention to how neurons in STN responded during beta bursts. STN neurons fired rhythmically in the beta frequency range (see autocorrelograms of example neurons during cortical beta bursts, black trace, Figure 3C). This firing pattern was more likely to occur for spikes that occurred during beta bursts than those that occurred outside beta bursts (t-test comparing power spectrum of neural spike trains that occur during vs. outside cortical beta bursts: STN beta bursts, t(82)=2.52, p=0.014; motor cortex beta burst: t(82)=2.93, p=0.0044) (Figure 3D, top plot). This suggests that STN neural beta-firing is linked to beta oscillatory activity in STN and motor cortex. Indeed, neurons were more likely to exhibit significant spike-field coherence with motor cortex than with sensory cortex (Supplementary Figure 4), indicating that STN beta might be more related to the preparation to execute a movment than sensory feedback from a movment.

21/83 (25%) neurons showed significant beta peaks in their autocorrelogram (see methods; Raz et al., 2000; Amirnovin et al, 2004). Both movement and stop-signal neurons had a propensity to coordinate their firing with beta rhythms (7/32 movement neurons fired at beta (22%); 2/10 stop neurons fired at beta (20%)). However, only movement neurons were more likely to fire at a beta rhythm during motor cortical beta bursts (Figure 3D bottom plot; t-test comparing average beta spike-train power during vs. outside beta bursts: movement neurons: t(31)=2.9, p=0.006; stop-signal neurons t(9)=1.56, p=0.15; non-responsive neurons: t(47)=1.18,p=0.24). This result suggests that motor cortical bursts are specifically linked to beta firing in movement neurons.

Previous studies (e.g., deHemptinne 2013; de Hemptinne 2015) suggest that exaggerated beta-synchrony between STN and motor cortex due to pathology may inhibit movement. If this is the case, we would expect neurons in STN to be more likely to fire at a beta rhythm at rest, when movement is not occurring. We calculated the beta firing in 1 second bins throughout the course of the task and found that coordination of firing with beta is strongest at rest and is transiently reduced during movement. For example, the neuron in Figure 3E exhibited a peak in its spike-train power-spectrum in the 13-30 Hz beta band, but this peak was reduced following target onset. In particular, on successful stop-trials (Figure 3E, red trace), this neuron reduced its beta-related firing less compared to latency matched go trials (dark blue trace) and to failed stop trials (orange trace). To quantify this effect, we averaged the activity across the beta firing neurons that met criteria for stop-signal analysis (n=10 neurons). Beta firing appeared to decrease earlier on fast go-trials compared to stop-trials, but this difference was not significant (Figure 3F, top plot; t-test comparing beta spike-train power 1 second window after target onset: t(9)=1.5, p=0.167). On successful stop trials, however, beta firing was significantly higher than on latency matched slow go-trials (Figure 3F, bottom plot; t-test comparing beta power 1 second window after target onset: t(9)=1.5, p=0.167). This suggest that the firing activity of STN neurons is strongly coordinated to ongoing beta activity both during rest and during successful stopping. This data supports the hypothesis that for a movement to be initiated, STN neurons must break out of beta entrainment to dynamically respond and execute a movement. This data also suggests that on successful stop-trials beta entrainment is less likely to attenuate following the target (Fig. 3F, bottom), an observation at odds with beta burst activity which shows a complete attenuation following the go-signal (Figure 3A). Beta-firing of single neurons, thus, may not necessarily reflect a 1-to-1 relationship with the local field potential, which represents a broader or more widespread view of ongoing brain activity.

### Features of neural spike trains predict recording location in STN

Stop-signal and movement neurons were located in different parts of the STN along the dorsal-ventral axis (Figure 2I). However, this result is based on the average location of these two groups of neurons, leaving open the question of whether knowing the properties of an individual neuron is sufficient to identify its location. We thus next evaluated whether the response properties and spike train features of a neuron would allow a decoder to determine the anatomical location of an STN neuron. We used 5 spike train features (firing rate, coefficient of variation, local variation, burst Index, beta-firing, and spike waveform), 2 spike-field features (local PPC between STN spikes and STN beta, PPC between STN spikes and motor cortex beta), and 4 task features (change in firing rate during movement, following stop signals, during successful stop trials, and during failed stop trials measured as an effect size). We then used classifiers to determine if these features predicted recording location.

First, in an unsupervised approach, we used principal component analysis to reduce the dimensionality of the feature space and then determined if any of the principal components described an anatomical axis through STN. The first 4 principal components (PCs) explained 71% of the variance in the features (Figure 4A). Through varimax feature rotation, we maximized the loading of each feature onto one of the four components (Figure 4A). Feature weights show that spike-train attributes primarily map onto components 1 and 4 (component 1= rate, LV and CV; component 4=burstiness and waveform), while task related features primarily map onto components 2 and 3. Component 3 is composed of movement features and spike-field synchrony, while component 2 is composed of stop-related features (Figure 4A). Unsupervised PCA analysis thus identified several axes that were primarily related to tuning of cells to either the stop signal or movement onset, whereas others were primarily related to firing statistics.

**Figure 4:**
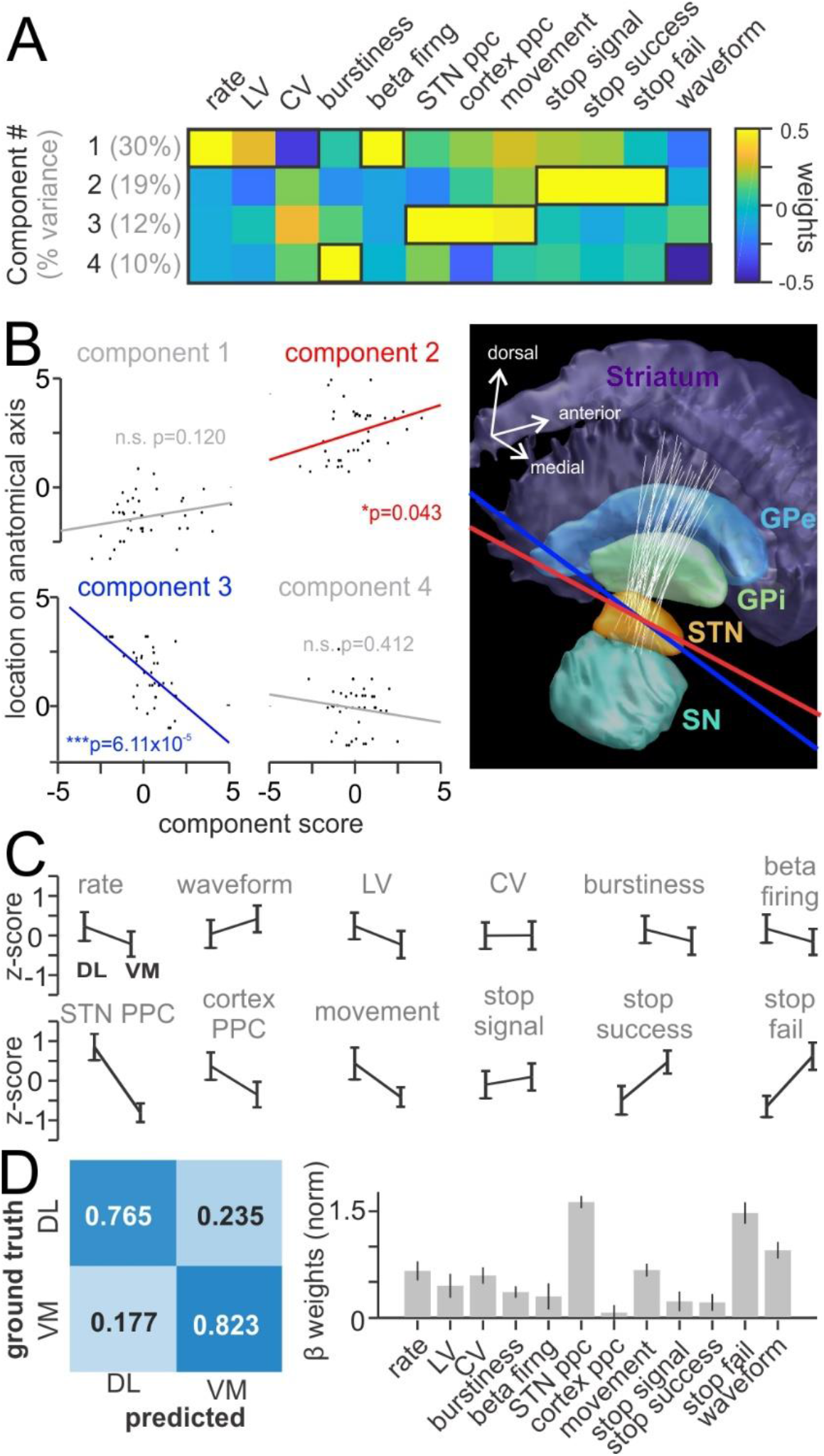
Spike-train characteristics of STN neurons vary along a dorsolateral-ventromedial anatomical axes through the STN. A. PCA applied to single unit firing features reveal four components/factors that together describe 71% of the variance in the data. B. Correlation of the scores of each component with the position along different anatomical axes through the STN. The scores of component 2 and 3 significantly correlated with the position along two axes (red and blue, respectively) that both spanned the dorsolateral-ventromedial range of the STN. Components 1 and 4 were not significantly correlated with any axes in STN (linear regression, significant slope coefficient, p<0.05). The illustration shows the axes that correlate with components 2 and 3 (red and blue lines) in anatomical space. White lines indicate the trajectory of each microelectrode. C. Average feature value for each feature projected onto a single dorsolateral-ventromedial axis through the STN (average of blue and red axis shown in (B)). Neurons were grouped onto either the dorsolateral region (DL) or ventromedial (VM) part of the STN by median split. Shown are the average±SEM spike train features for DL and VM neurons (rate=mean firing rate; waveform=narrow or wide; LV=local variation; CV=coefficient of variation; burstiness=burst index based on autocorrelogram; beta firing=power of spike-train in beta range; move, stop-signal, stop-success, and stop-fail= effect sizes of neural firing rate changes during the task) D. (left) Performance (confusion matrix) of an SVM classifier trained to discriminate DL from VM locations of neurons based on (C) tested on left out neurons. (right) The average feature weights of the SVM classifier. Higher weights indicate that this feature plays a greater role for classification. *p<0.05, **p<0.01, ***p<0.001

To identify which, if any, of these four PC axes describe anatomical space in STN we correlated the component scores of each neuron with the location of that neuron projected onto different anatomical axes through STN (see methods). This procedure revealed two axes that significantly correlated with components 2 and 3, the task related components (significance assessed by linear regression, p<0.05; components 1 and 4 did not significantly correlate with any axes) (Figure 4B). These axes spanned the STN largely along a dorsolateral-to-ventromedial axis (red and blue lines in Figure 4B). While the scores of the movement component (PC3) decreased moving dorsolateral-to-ventromedial along this axis, the scores of the stop component (PC2) increased (Fig. 4B). Intriguingly, these axes, which best differentiate stop and movement activity, run approximately perpendicular to the tracts of the recording electrodes (white lines, Figure 4B anatomical reconstruction). This indicates that the depth along the recording track is not a necessarily a good differentiator between these two types of STN functions.

The two anatomical axes we identified with PCA follow a similar dorsolateral-ventromedial axis. To simplify analysis, we averaged the position of these two axes to yield a single axis through STN that describes both components. We then projected the location of each STN neuron onto this axis and identified whether it existed in the dorsolateral region of the axis or the ventromedial (median split of all neuron locations). Many of the neural features differed between these two anatomical locations (Figure 4C, e.g., stop-related features have higher values in the ventral region while movement-related features are higher in the dorsal region), indicating that this summary axis is an appropriate way to parcellate the data.

Lastly, to see if the firing features of a neuron predicted which region of the STN a neuron was recorded from, we trained a classifier to differentiate the dorsolateral region from the ventromedial region using the firing features of the neurons. We tested the performance of this classifier on neurons not included in training and achieved a performance of 79% (Figure 4D; chance=50±11%, t(999)=86.7, p=3.6 x 10^−467^). The weights assigned by the classifier to different features indicated that a variety of features contributed to classification, especially spike-field features and task-features (Figure 4D). Indeed, if only spike-field and task-features were used to train the classifier, the classifier performed significantly above chance at 71% accuracy (Supplementary Figure 6; chance=50±11% by shuffling labels 1,000 times; t-test comparing performance to shuffled values: t(999)=63.3, p=6.9 x 10^−352^). Importantly the classifier was specific to the optimal axes through the STN. If a different axis was used, for example a dorsal-ventral MNI axis aligned to anatomical midline and the anterior/posterior commissure, the classifier performed at chance (Supplementary Figure 7).

Together, these set of analysis strongly suggest that neuronal firing features describe anatomical subfields in STN with distinct functional roles. While the dorsolateral region exhibited neural responses indicative of a go pathway through STN, the ventromedial region exhibited responses representative of a stop pathway.

### Parkinsonian state

Throughout this paper we have presented data on STN activity in humans, all of whom have Parkinson Disease. Because we cannot record single neuron data from STN in healthy controls, it’s difficult to differentiate effects that are due to pathology and those that are part of normal brain function. To attempt to identify neural activity due to pathology, we separated the data into groups of patients with low vs. high UPDRS III scores, a routine neurological test of the motor symptoms of Parkinson Disease (median split; median=31, low UPDRS group: 21±8, high UPDRS group: 47±16).

Patients with low and high UPDRS scored performed similarly on the task. They did not differ in their behavioral response times (low= 957±341 ms, high=900±322 ms, two sample t-test comparing response latencies t(17)=0.369, p=0.717), their SSRTs (low: 438±366; high: 410±338; t(14)=0.16, p=0.875), or their response accuracy (movement in correct direction instructed by go-signal: low= 95±6%; high: 97±2%; t(17)=1.05, p=0.308; successful stopping accuracy: low= 44±2%, high=46±2%; t(17)=0.177, p=0.862);

There was an equal likelihood of recording movement neurons in patients with low or high UPDRS scores (P(movement neuron)= low: 0.35±0.20; high= 0.41±0.22; t(14)=0.441, p=0.67), as well as stop-signal neurons (P(stop signal neuron)=0.17±0.26; 0.19±0.27; t(11)=0.06, p=0.95). None of the firing rate features of the neurons, the spike-field beta synchrony, or the task responsiveness significantly varied among the two groups (Supplementary Figure 8). Together, this indicates that our results did not differ systematically with Parkinson Disease symptom severity as assessed by UPDRS III, suggesting that the patterns of activity we found were not due to Parkinson Disease.

## Discussion

The discovery of hyperdirect pathways has sparked a breadth of new research positing a broad role of the STN in selecting (Zaghloul et al., 2012; Frank, 2006), canceling (Schmidt et al., 2013; Pasquerue & Turner, 2017; Aron & Poldrak, 2006; Chen et al., 2020), switching (Isoda & Hikosaka, 2008; Pasquereu and Turner, 2017; Jantz et al., 2017), and pausing actions (Fife et al., 2017; Dutra et al., 2018) through activation of a stop pathway. Here, we show that the human STN participates in selecting and cancelling movements at the level of single neurons. The properties of these neurons vary systematically along the dorsolateral-ventromedial axis of the STN, which is different from the previously hypothesized but never experimentally verified dorsal-ventral axis of neuronal tuning with respect to movement vs. stop signals. We further show that stop-related activity occurs sufficiently quickly to play an active role in stopping, whereas the typically hypothesized increases in beta activity do not.

Race models of action cancellation propose that activity in a “stop-pathway” rapidly increases after encountering a stop-signal (Boucher et al., 2007; Schmidt et al., 2013; Wiecki and Frank, 2013). Here we identified neurons in the STN whose response properties indicate that they are part of this stop pathway. Stop-signal neurons were activated shortly after the appearance of the stop-signal regardless of behavioral outcome (success or failure to stop). Similar neurons have been observed in the STN of mice (Schmidt et al., 2013) and macaques (Pasquereu and Turner, 2017), but have so far not been shown in humans. This across-species commonality suggests a shared mechanism of stopping, a fact supported by the observation that different species exhibit similar behaviors during the stop-signal task (Middlebrooks and Schall, 2014; Kornylo et al., 2003). Neurons that are activated by stop-signals have been previously reported in humans (Benis et al. 2016; Bastin et al. 2014). However, these cells were only activated by successful (and not failed) stop trials, a profile of response not compatible with neurons in the stop pathway. In contrast, in our data, only 2% of neurons met these criteria of only being activated by successful stop trials; a larger portion (11%) registered the appearance of a stop signal regardless of behavior. Note that while we recorded relatively few stop-signal cells (10/83), this proportion is not unexpected given the tendency for recordings in more dorsal STN (where such neurons are more rare) and is similar to the proportion of such cells observed in macaque STN (n=14 stop cells in monkeys, 8% of the recorded population in the study of (Pasquereu & Turner, 2017). Together with this prior study, our data indicates that these stop-neurons are rare compared to other types of STN activity.

While neurophysiological models of action cancellation often emphasize the role of the STN in the “stop pathway” (Wiecki and Frank, 2013; Schmidt et al, 2013), here we also observed a large number of neurons involved in movement. These neurons are part of the ‘go pathway’ as suggested by studies in humans and monkeys (Lipski et al., 2018; Potter-Nerger et al., 2017; Pasquereu & Turner, 2017; Georgopolous et al., 1983). Most of these movement-related neurons *began to fire* prior to movement onset, which supports their role in the “go-pathway” that selects/executes actions. The majority of these neurons didn’t achieve peak firing rates until after the movement started. From the point of view of a threshold-model of action selection/execution, this pattern of activity is unexpected. However, most threshold-models are based on a ballistic movement, i.e. a saccadic eye-movement or button press. In contrast, here our subjects used a joystick to respond, which likely requires continual sensory feedback and updating of the ongoing motor plan. Indeed, on some failed stop trials, subjects adjusted their movement and returned the joystick to the center position before reaching the full extent of the response. The early response of movement-related neurons might thus reflect a role in action selection while later responses might reflect proprioceptive feedback and/or updating of the motor plan. While classical models of the basal ganglia segregate “direct” go-pathways from “hyperdirect” and “indirect” stop-pathways, more recent models suggest that the three pathways are heavily intertwined (Calebresi et al., 2014). Our data support the view that “stop” and “go pathways” in the STN are intermixed.

Some of the movement-related neurons we reported reached their peak firing rate at the time of movement onset as predicted by a threshold model of action execution. These neurons are reminiscent of neurons in motor and decision making circuits that exhibit ramp-like increases in activity until a motor or decision-threshold is reached (Herz et al., 2016; Hanes and Schall, 1996; Schall, 2019). Indeed, we found that these neurons achieved similar peak firing rates regardless of response latency (a characteristic of threshold like responses) but they ramped to this peak firing rate faster on fast reaction time trials. As would be expected of an interactive race-model of action cancellation (Boucher et al., 2007), where the go-pathway interacts with the stop-pathway, movement-related neurons attenuated their firing rate after stop-signals if the action was cancelled successfully.

How can an increase in firing rate in the STN, which is a nucleus with exclusively excitatory output (Smith and Parent, 1988; (albeit it does contain inhibitory interneurons Levesque and Parent, 2005), cause two seemingly disparate actions: moving and stopping? One possibility is that movement and stopping neurons in STN project to two different targets, e.g., stop-related neurons might directly excite SNr or GPi to inhibit movement, while movement-related neurons might excite GPe which in turn inhibits SNr and GPi to facilitate movement (de Vito et al., 1980; Parent and Smith, 1987; Joel and Weiner, 1997; Jantz et al., 2017). Another possibility is that movement and stop-signal neurons both lead to an inhibition of motor programs. In this vein, activity of movement-neurons doesn’t reflect the excitation of desired motor program, but instead reflects inhibition of competing motor programs (e.g. as hypothesized by a center-surround mechanism of action cancellation, Mink 1996). This inhibition helps facilitate the execution of the desired program. When stop-signal neurons fire, they might first (1) globally inhibit all motor programs (including the desired one) (Wessel et al., 2016) and (2) attenuate the response of movement-neurons in STN to reduce the level of surround inhibition. Compatible with this view, we found that movement-related neurons in STN tended to be non-selective to movement direction (left/right; muscle adduction/abduction).

Tract tracing studies in monkeys have revealed that different parts of the STN receive hyperdirect input from distinct cortical regions (Haynes and Haber, 2013). A tripartite model divides STN into motor, associative, and limbic regions that are thought to play distinct roles in action selection, cognition, and emotion (Greenhouse et al., 2011, 2013; Jahanshai et al. 2015ab; Mallet et al., 2007; Ewert et al., 2018). Our data provides support at the single-neuron level for such a parcellation in humans. In line with other studies of STN neural activity (Starr et al., 2003; Theosopoulous et al., 2003), movement-related neurons predominantly clustered in dorsal STN, the primary target for treating motor symptoms with DBS therapy. Stop-signal neurons were localized more ventrally, similar to observations in macaque (Pasquerau and Turner, 2017). The right inferior frontal gyrus, the most likely cortical candidate for cancelling actions (Aron et al., 2016), is thought to terminate in STN more ventrally and is activated antidromically by electrical stimulation of the STN (Chen et al., 2020; Haynes and Haber, 2013).

What are the clinical implications for our results? Single unit studies in STN have suggested that movement-related firing features (Starr et al., 2003), oscillatory firing patterns (Shimamoto et al., 2013, Fischer et al, 2020), or bursty firing features (Seifried et al, 2012; Kaku et al. 2019) vary between subfields of the STN and can therefore be used to guide placement of DBS electrodes that provide therapy for motor disorders. We developed a classifier that predicted the recording location of a single neuron based on firing features and task parameters. While movement- and stop-related responses had predictive value, even non-task related features significantly contributed to the prediction of dorsolateral-ventromedial location. These features could thus be used to help better target electrodes during surgery and focus stimulation (e.g. Zhang et al 2020). Intriguingly, even the location of neurons in the most ventromedial region of STN (which were non-responsive during the stop-signal task) could be accurately decoded. Based on a hyperdirect parcellation of STN, these neurons might represent the limbic domain within STN. Future experiments are needed to determine whether these neurons are activated by tasks that involve limbic areas such as error monitoring, reward valuation, emotion, and anxiety (Wagenbreth et al., 2019; Greenhouse et al., 2014; Gourisanker et al., 2018; Fu et al, 2019). More broadly, from a clinical perspective, the anatomical variation we have shown along the dorsolateral-ventromedial direction may explain why stimulation in ventral STN improves tremor in Parkinson Disease but also results in a side effect of more impulsive behaviors (Ballanger et al., 2009; Voon et al., 2017; Hershey et al., 2010).

We did not observe significant differences in the behavior or the single neuron activity in patients with high vs. low UPDRS III scores. While at first surprising, other studies have shown that DBS, which typically lowers UPDRS scores, can lead to widely different patterns of reaction times and SSRTs in the stop signal task (Ray et al., 2009; van den Wildenberg, 2006). UPDRS III scores may not be the optimal descriptor for differentiating Parkinsonian severity in our sample, as these scores are broad and include many aspects of motor dysfunction. A single UPDRS III score does not necessarily differentiate the dominant motor features of disease in each induvial such as bradykinesia, tremor, or rigidity, and it is possible that further sub-analysis of these features may reveal some correlations to neuronal activity recorded. Nonetheless, given the fluctuating nature of Parkinson disease, the variable use of medications by patients before surgery, and similar variables, the UPDRS III score provides a reasonable single metric to suggest that the underlying disease itself had relatively little impact on our findings. Future experiments that repeat these studies in a different patient population (e.g. cervical dystonia) are needed to reveal the extent to which these firing properties vary among different subject pools.

Last, similar to other studies, we observed that neurons in the STN have a high propensity to spike at certain phases of the beta rhythm, especially in patients with Parkinson Disease (Shimamoto et al., 2013; Yang et al., 2014; Kuhn et al., 2005; Lipski et al., 2017). While an increase in beta activity at the level of the local field potential is often assumed to be the mechanism by which stopping is implemented (Alegre et al., 2013; Bastin et al., 2014; Wessel et al., 2016a; Fischer et al., 2017, Ray et al., 2012), this prediction has yet to be validated at the level of single neurons. Contrary to this prediction, we found that while beta activity increases following successful stopping, this increase appears too late to be able to contribute to successful stopping. Therefore, we conclude that the elevated beta activity seen during successful stopping reflects either the termination of the stop process or maintenance of the stop process rather than the detection of the stop signal itself. Instead, we show here that only the stop signal at the level of single neurons is fast enough to be able to initiate stopping.

## Methods

### Subject Details

Electrophysiological signals were recorded intraoperatively from patients with Parkinson Disease undergoing surgery for the implantation of a Deep Brain Stimulating device. A total of 19 subjects (33 sessions/surgeries) volunteered to participate in this study (see Tables S1 for demographics, age at time of recording = 68±7 years, UPDRS III score off medication, pre-DBS treatment: 33±18, n=8 females). Subjects often performed two sessions of the task, during surgery for each side of the brain leading to 33 total sessions. 5 sessions were excluded from analysis because the patient was unable to complete the task (n=1) or we were unable to isolate stable neural recordings (n=4), leaving 28 sessions for analysis. Subjects were off their Parkinson medication for at least 12 hours prior to surgery. All subjects gave informed consent and all protocols were approved by the Institutional Review Board of Cedars-Sinai Medical Center.

## Method Details

### Task

Subjects performed a stop-signal task (Aron & Poldrack, 2006; Verbruggen et al., 2019). On each trial the go-signal (white arrows) appeared on a computer monitor instructing the subject to move a joystick (Thrustmaster T16000M, Thrustmaster, Hilsboro OR) left or right (Figure 1A). Subjects performed the task with the hand contralateral to the recording site. On a random subset of 33% of trials a stop signal appeared (red arrows) following the go signal at a variable delay. The stop signal delay (SSD) was initially set to 300 ms and was then automatically titrated using an adaptive staircase method to achieve a ^~^50% rate of successful stopping. If a subject was successful on a stop trial, the SSD was reduced by 50 ms on the subsequent stop trial; if they failed to stop the SSD was increased by 50 ms. During the experimental recording, a trial was registered as a movement if it exceeded a pre-defined threshold (20 degrees of the total extent of the joystick). On go trials, the trial turned to a blank screen after 1.5 seconds or after a movement was detected, whichever came first. On stop trials, the trial was terminated after 1.5 seconds if no movement was detected. At the termination of a trial, the screen went blank and was followed by a 1 s intertrial interval. No trial-by-trial feedback was provided. Subjects were trained to perform the task pre-operatively (20-30 trials). They were instructed to “move the joystick in the direction of the white arrows as quickly as possible and, if the arrows turned red, to treat it like a stop-light and attempt to stop performing the movement.” Subjects performed 120 trials during surgery (40 stop trials total, a break was given midway through the session). The task was implemented in MatLab (Mathworks, Natick MA) using Psychtoolbox (Brainard, 1997 http://psychtoolbox.org/;).

### Electrophysiology

Micro-electrode recordings of single neuron activity were performed during anatomical mapping of the STN during surgical implantation of a Deep Brain Stimulating device (for surgical details and target mapping see Kaminski et al, 2018). Micro-electrodes (Alpha Omega Hybrid microelectrodes, STR-000080-0024950) contained a micro-wire at the tip and a larger macro-contact 3 mm above the tip. Throughout the paper we analyze the single unit activity and local field potentials from the micro-wire. In each session, 1 microelectrode was placed along the target trajectory and a second electrode was placed in a Bengun array either 2 mm medial/lateral or anterior/posterior. Electrodes were grounded and referenced to the guide cannulae placed 25 mm above the target site. The broadband 2 Hz-44 kHz signal was recorded using an NeuroOmega system (Alpha Omega Inc.). The two microelectrodes were advanced simultaneously in 1 mm steps while monitoring single neuron activity and comparing it to the expected trajectory determined by surgical imaging. Upon approach to the STN, the speed of the electrode was reduced to 0.1-0.5 mm/step and single neurons were monitored. When we detected single unit spiking on at least one of the micro-electrodes we waited at least one minute to ensure the spike amplitude remained stable before beginning the experiment. Following the experiment, the electrode was further advanced below target to identify the location of the substantia nigra pars reticulate (SNr), a lower boundary of STN. We used electrical stimulation of SNr (30 uAmp train at 200 Hz), which reliably leads to a pause in firing rate to further confirm the lower boundary of the STN (see Lafreniere-Roula et al., 2009; Kaminski et al, 2018)

An ECog strip (AdTech IS08RSP10X-0T1, 8 contacts, 10mm contact spacing,) was inserted subdurally as published previously (Kaminski et al., 2018) to record intracranial EEG signals over sensorimotor cortex. Ecog recordings were referenced to a needle placed in the skin over the mastoid process and grounded with a surface electrode at the same location. iEEG signals were recorded with a 2-350 Hz bandpass filter, simultaneously with the recordings obtained from the mircoelectrode. We refer to recordings made from the subdural strip electrodes as iEEG.

### Electrode localization

Recording locations of single neurons were determined by fusing pre-surgical MRIs with a post-surgical CT scans to visualize the location of the DBS electrode. First, the post-surgical CT scan was co-registered with the pre-operative MRI scan (6DOF transformation using BRAINS registration, 3D slicer. Federov et al., 2012). The tip of the DBS electrode was marked manually on the CT-scan (last CT trace with visible contrast) and traced along each slice of the axial CT. This generated a cloud of points that marked the trajectory of the DBS electrode. We fit a line to this cloud of points by using principal component analysis: PCA was applied to the marked locations, the marked locations were projected onto the first principal component to identify the main axis, and we constructed a normal vector along this axis with its origin at the electrode tip. This normal vector describes the axis of the DBS electrode. The recording electrode is along the same trajectory as this normal vector but may be driven to a different depth or placed at a different medial-lateral or anterior-posterior location in the ben gun array (notes were made during surgery to record this offset). To determine the medial-lateral position we calculated a second normal vector that was orthogonal to both the DBS electrode vector and to the plane that defines patient midline. A third normal vector that was orthogonal to both these vectors described the anterior-posterior position and completed the basis used to determine the microelectrode recording sites. The position of a neuron in electrode space (e.g. 1 mm deep relative to the DBS electrode and 2 mm lateral) is multiplied by this basis to determine its position in native patient space. To co-register data across patients, we used an MNI coordinate system. The patients MRI was aligned to the 2009b NLIN Asym ICBM template brain (Fonov et al., 2009) using an affine transformation followed by a symmetric image normalization (SyN) diffeomorphic transform (Freesurfer, http://surfer.nmr.mgh.harvard.edu/). We then applied these transforms to the neuron coordinates in native patient space to obtain the coordinates in MNI space. Locations of neurons are plotted together with the DISTAL atlas specialized for subcortical structures targeted in DBS surgery (Ewert et al, 2017).

During the recordings, iEEGs from the ECog strip were referenced to an electrode placed in the skin near the mastoid process. The location of the electrodes in the brain was determined by identifying a reversal in the polarity of the evoked potential in the iEEG caused by stimulation of the median nerve of the contralateral hand (20 – 30 mA, Cadwell Elite system) (Allison, 1987). This method (Allison et al., 1987, Kaminski et al, 2018, de Hemptinne et al., 2013) has previously been used to reliably identify the orientation of contacts anterior to and posterior to the central sulcus. For analysis, we performed bipolar referencing by calculating the difference in the signal recorded by adjacent pairs of contacts. We refer to the bipolar pair where the reversal took place as the central sulcus (CS) (e.g. if the reversal occurred between electrodes 2 and 3 during the recording, as in Figure 1 H, the bipolar pair 2-3 was labeled CS). The pair immediately anterior to the CS was identified as motor cortex.

## Quantification and Statistical Analysis

### Behavior

Trials were categorized as “successful stop” trials if the joystick remained within the detection threshold for at least 1.5 seconds. Stop trials where movement was detected were considered “failed stop trials”. During our analysis we noticed that sometimes subjects moved the joystick on stop trials but returned the joystick to center before reaching detection threshold (20 degrees). We decided that these trials with a small amount of movement should be considered failed stop trials post-hoc for analysis purposes. For analysis we thus set the threshold to a lower value (11 degrees) based on the distribution of joystick values at trial start time (99.5% of the horizontal joystick positions are <11 at rest during the first 250 ms following the go-signal). On go-trials and failed-stop trials, reaction times were measured as the point of time after the go-signal when the joystick position first significantly exceeded its position at rest (two standard deviations).

In each session we calculated the percent of stop trials in which the subject successfully cancelled the action (i.e. did not move past the 11 degree threshold). We calculated the stop-signal reaction (SSRT) time for each session using the integration method (Verbruggen et al., 2019). Briefly, this method involves first calculating the cumulative distribution function for the reaction times across all trials (including all go trials and failed stop trials) and then identified the latency L at which P(latency<L)= P(stop success rate | stop trial). Go-trials with reaction times <L are considered fast go-trials. Thus, if P(stop success rate | stop trial)=0.5, go-trials are split into fast and slow-go trials as a median split. According to the integration method, SSRT = L – mean(SSD on stop trials). We calculated the inhibition function for each session by dividing trials into those with short and long SSDs (median split) and calculating the percent of failed-stop trials for each. For stop-signal behavioral and neural analysis we only used sessions where the subject performed at least 5 successful and 5 failed stop trials. For stop-related analysis we calculated fast- and slow-go trials using the integration method described above. For movement-related analysis (Figure 2A-D) we defined fast and slow-go trials using a median split in reaction time to maximize the number of neurons available for analysis.

### Spike detection and sorting

Broadband signals were filtered in the 300-3,000 Hz band using a zero-phase lag filter (four-pole Butterworth filter). Spikes were detected and sorted using the semi-automated template-matching algorithm OSort (Rutishauser et al., 2006). Each single unit was manually evaluated and verified based on the following features: (1) the spike waveform, (2) the percentage of interspike intervals (ISIs) less than 3 ms, (3) the ratio of the waveform extremum and the standard deviation of the noise, (4) the pairwise projection distance in the clustering space between all isolated neurons on the same microwire, and (5) the coefficient of variation of the ISI and (6) the cluster isolation distance (Rutishauser et al., 2008) (spike quality metrics, Supplementary Figure 1). To account for instabilities in neural recordings, we manually selected a lower and upper time limit during which a neuron achieved firing stability and only analyzed these trials. 26 neurons (31%) were stable for all 120 trials of the recording (15-20 minutes), whereas the remainder was stable only for a subset of trials. On average, the neurons were stable for 89±21% (89 ±37 trials) of the recording.

### Spike waveform analysis

We calculated the average spike waveform of each neuron and quantified three width features of the waveforms: the trough-to-peak width (time from the spike trough to the next peak, corresponding to the after hyperpolarization), the half-width (the width of the spike at half the spikes amplitude) and the repolarization time (the time elapsed from the peak hyperpolarization to the time this peak is at its half amplitude) (for detailed methods on how these widths are calculated see Mosher et al., 2020). Plotting these three width features revealed two clusters of neurons, a narrow and a broad spiking group (Supplementary Figure 5). We used a published classifier that differentiates between narrow and broad spiking neurons (Mosher et al., 2020) to determine if a neuron belonged to a narrow or broad cluster.

### Identification and analysis of movement-related neurons

A neuron was classified as “movement-related” if it exhibited a difference in firing rate during movement compared to a pre-target baseline. On each trial, firing rate was calculated in a 200 ms sliding window with 1 ms overlap. A non-parametric Wilcoxon rank sum test compared each bin during movement (−250 to 500 ms aligned to movement onset) to all bins at baseline (−500 ms to 0 ms before target). This identified bins during movement with potentially significant differences in firing rate from baseline (p<0.05). To correct for multiple comparisons, we performed a cluster-correction analysis (Maris and Oostenveld, 2007). In this procedure a cluster is defined as a set of adjacent time bins with potentially significant firing rates. We sum the test-statistic (rank sum z-value) across the bins in a given cluster to give a “cluster-wise statistic”. We repeated the analysis 500 times but randomly shuffled bins that occur during baseline and movement. If the cluster-wise test-statistic of the data exceeded 95% of the cluster-wise values obtained from shuffled data, the neuron was said to be significantly movement responsive.

In addition to aligning the neural spike times to movement onset, we aligned spikes to the onset of the go-signal (Figure 2B). We determined if the response of a neuron was more tightly locked to movement or the appearance of the go-signal by identifying the peak firing rate (200 ms binned PSTHs, 1 ms sliding window) after aligning to either time point and then compared the peak firing rate in the two conditions with a paired t-tests (Figure 2B inset).

Movement neurons had a ramp like response in firing rate leading up to movement onset. To measure this phenomenon, we binned firing rate (200 ms bins, 1 ms overlap) and fit a line using least squares regression to these rates from the timepoint when the neuron started to change its firing rate (cluster method, see above) to the timepoint of movement onset. Neurons with firing rate onsets that began after movement onset were excluded from this analysis.

We determined whether the population activity across all recorded neurons differed between periods of movement and periods when the subject was stationary sufficiently strongly to allow single-trial decoding using a decoder. We based this analysis on a pseudopopulation. First, we set aside one trial of each type (fast go, slow go, success stop, failed stop) for each neuron. These trials were later used for testing. We then calculated the mean firing rate on every trial during movement (1000 ms window following movement onset) and rest (1000 ms window, beginning 1500 ms before movement onset). For each neuron, we normalized the firing rates (z-score) across all trials. We randomly selected 60 trials from each neuron and created an n x m x 2 matrix of firing rates (n=neuron number, m=trial, 2=rest or movement period). We trained a linear SVM classifier on these firing rates to decode rest from movement and tested this classifier to the left out trials. We repeated this entire procedure 1,000 times. To examine the time course of this decoder throughout the trial, we binned the firing rates of the left-out trial data (200 ms bins, 50 ms overlap) and applied the decoder to each bin. The trace in Figure 2G shows the probability that the decoder predicts movement correctly as a function of time. As a control, we repeated this entire procedure 1,000 times but shuffled the rest and movement labels across trials. This generated a distribution of performance values expected by chance. To assess significance, we subtracted the performance value of the true decoder from the performance of each shuffled decoder. If the distribution was significantly different from 0 (t-test), the decoder was said to perform significantly better than shuffled data.

### Identification and analysis of stop-signal responsive neurons

A neuron was classified as “stop-signal responsive” if it’s firing rate following the onset of a stop-signal was significantly different from the pre-stop baseline. We used the same binning, test-statistics, and clustering procedure as we did for selecting movement neurons (see above, rank sum tests, cluster-correction technique), but instead aligned to onset of the stop-signal. A cell was considered stop signal responsive if any bin in the 1000 ms following a stop-signal was significantly different from the firing rates observed in the window 500 ms immediately preceding the stop-signal. Cells for which less than 5 successful or failed stop trials were available were not included in this analysis.

The above selection criteria does not require stop-signal responsive neurons to differentiate between trial outcome (success or failure to stop). To determine if neurons responded to stop-signals regardless of trial outcome we in addition applied a stricter criterion with two statistical tests. First we compared the binned firing rate on successful stop trials to latency matched slow go trials (same 200 ms bins, statistical test is Wilcoxon rank sum as used previously for classifying stop-signal responsive and movement neurons, go trials were aligned to the time when the stop-signal would have appeared). Second, we compared the binned firing rate on failed stop trials to fast go trials. If the firing rate in any bin within 1000 ms after stop-signal onset significantly differentiated successful stop trials from slow go trials and failed stop trials from fast go trials, the cell was said to have met this strict stop-signal criterion.

For each neuron we calculated the effect size of the stop-signal on neural firing (Cohen’s D). We compared the effect of the stop-signal relative to baseline (mean rate in 500 ms window before stop signal compared to 500 ms window after stop signal). We also compared the effect of successful stop vs. slow go trials and failed stop vs. fast-go trials (rate 500 ms after stop signal onset, or the time when the stop signal would have appeared on a go-trial).

To better understand how the population of neurons in the STN might detect stop-signals we trained single-trial population decoders, similar to the movement decoder described above. We first calculated the firing rate in 200 ms bins with a 50 ms sliding window for every trial and every neuron. Then, like with the movement decoder, we randomly set aside one trial from each trial type (fast go, slow go, successful stop, failed stop) for later testing. We next randomly selected 30 stop-signal and 30 go-signal trials. In the movement decoder, we trained using the firing rate at different time points within the *same* trial (movement vs. pre-movement period). In the stop-signal decoder we compare across *different* trial types. For each neuron, we normalized the firing rate at a given time (z-score, normalized across both go and stop-signal trials). We then trained a linear SVM classifier on these normalized firing rates at a given time bin and tested on all time bins for left-out trial. We repeated this train and test procedure for all bins spanning 1,000 ms before to 2,000 ms after the stop signal. This generated a matrix of decoder performance for a single pseudopopulation of neurons. We repeated this procedure 1,000 times with different left out data and different training subsets and robust composite classifiers each composed of 1,000 classifiers trained on different subsets of data and tested on left-out data. To assess whether any of these classifiers performed better than chance, we repeated the entire procedure on data with shuffled labels for stop and go-signal trials (1,000 shuffled bootstrap). To determine if the real data was significantly different from the shuffled data we z-scored the performance of the classifiers to these bootstrapped values and detected every time bin that performed better than chance (p<0.05). We applied a 2D cluster analysis to account for multiple comparisons. First, we identified adjacent time points in 2d space (train x test matrix) with significant z-scores and summed their z-scores to give a cluster-wise statistic. We repeated the procedure on the 1,000 shuffled decoders, identifying all clusters and summing the values. If a clusterwise statistic exceeded 95% of the shuffled cluster statistics, the cluster was regarded as having a decoder performance significantly better than chance. The heatmap in Figure 2G shows the difference in decoder performance on true vs. shuffled data, contours outline significant clusters.

### Beta bursts and beta-firing rhythms

Beta bursts were detected using a thresholding method (as in Tinkhauser et al., 2017; Torricellos et al., 2018). We filtered the LFP/iEEG in the 13-30 Hz band using a zero-lag bandpass filter (Hamming, order=140, 3x the median beta frequency 21.5 Hz). We then applied the Hilbert transform to the signal and detected periods when the power exceeded a threshold equal to 75% of the distribution of all power values at all time-points during the task. A beta burst was detected if the power exceeded this threshold for at least 93 ms (2 cycles of a beta rhythm, median frequency 21.5 Hz). In Figure 3 we identify periods when no beta bursts occur as timepoints when beta power was less than 25% of the distribution of power during the task maintained for at least 93 ms.

To calculate spike-field coherence between STN spikes and beta oscillations we used pairwise phase-consistency (PPC) as a metric (Vinck et al., 2010). Field potentials (recorded from microelectrode tip on STN electrode or from contacts on Ecog electrode) were filtered in the 13-30 Hz frequency band. We then applied the Hilbert transform to the signal and calculated the phase of the signal. We then calculated the PPC using the phase at the time of the spike. We labeled a cell-FP pair as having significant PPC (Figure 4fg) if its PPC exceeded 95% of the PPC values for the shuffled data (LFP trace was shifted by a random value up to 5 seconds, 1,000 bootstrap shuffles).

For each neuron we calculated the autocorrelogram (5 ms bins, −300 to 300 ms centered on spike) and the power of the spike train during motor cortical beta bursts and in the absence of beta bursts (Thomson multi-taper power spectrum, time-bandwidth product 1.25, with the minimum frequency resolution of ^~^10.2 Hz given beta burst are defined to be at least two cycles or ^~^100 ms; pmtm function in Matlab). We standardized the power-spectrum to account for differences in power by dividing by the area under the power spectrum. In figure 3 we compare the standardized power spectrum during and outside beta bursts for all neurons (and the mean power in the 13-30 Hz band for different types of neurons. We identified neurons with significant peaks in the beta band using the method of Raz et al., 2000: the spike train power spectrum is first normalized by subtracting the mean power spectrum in the 5-40 Hz band and dividing by the standard deviation. Second, if the maximum standardized value in the beta band (3-13 Hz) exceeded 3 standard deviations then the cell was said to significantly fire at a beta rhythm (Raz et al., 2000; Amirnovin et al, 2004). We also compared the maximum power in the beta band (13-30 Hz) during vs outside beta bursts (Figure 3, t-tests). To compare between task conditions (Figure 3E,F) we calculated the power spectrum in a 1 second bin with a 100 ms sliding window.

### Latency of neural responses

The onset times of movement- and stop-signal responsive neurons were calculated as the first significant time bin (200 ms bin, 1 ms sliding window, Wilcoxon rank sum test) that survived clusterwise-correction (see above sections on quantification of movement and stop-signal responses).

The probability of a beta bursts increased after a stop-signal on successful stop trials. The onset of this beta activity was calculated as the first time point (1 ms resolution) where the probability of a beta burst was significantly different between successful stop and latency-matched slow-go trials (two-sample t-test, clusterwise correction).

To compare the response onset of stop-signal neurons to beta activity we also used a binless differential latency approach (Figure 3B, bottom plot). For each stop-signal neuron, we calculated the cumulative number of spikes elicited on each trial from 0-3 seconds after the stop-signal (1 ms resolution). We then calculated, at each time point, the difference in the average cumulative spiking on successful stop and slow go trials. We compared, at every timepoint, these difference across the population of stop-signal neurons (t-test) and calculated a cluster-wise test statistic (sum of all t-values that occur adjacent in time and p<0.05). We repeated this procedure 1,000 times with randomly shuffled labels to generate a null distribution of cluster-wise statistics. The response latency of the neuron was identified as the first timepoint whose cluster-wise statistic exceeded 95% of the shuffled data (Xiang & Brown, 1998; Rutishauser et al., 2015)

### Classification of STN recording location based on firing features

For each neuron we calculated several features of the spike train and used these features to predict recording locations. The feature space consisted of (1) mean firing rate, (2) CV2 (a metric of spike train variability, Nawrot et al., 2008), (3) burst index (the average value of the autocorr function in a window 3-20 ms after each spike), (4) the degree to which a neuron spikes at a beta rhythm (average value of the spike-train power spectrum in the 13-30 Hz frequency band), (5) the effect of movement on neural firing rate (comparison of mean firing rate during movement compared to pre-target baseline, measured as Cohen’s d), (6) the effect of stopping on neural firing rate (comparison of mean firing rate during successful stop trials and slow-go trials, measured as Cohen’s d), (7) the spike-field synchrony between STN neurons and local STN beta rhythm (PPC between STN spikes and beta-filtered LFP from STN, see above section on beta), and (8) the spike-field synchrony between STN neurons and motor cortex iEEG beta (PPC between STN and motor cortex). All values are normalized and reported as z-scores (Figure 6a). Note that not all features could be calculated for every neuron (e.g. in some recordings there were not a sufficient number of successful stop trials to calculate the stop effect size; some recordings did not have an SSEP reversal so we could not localize electrodes to motor cortex and calculate the motor related PPC). All features could be calculated for 37 neurons, which is the subset which we use for this analysis.

For the unsupervised classification, we performed principal component analysis on the firing feature space and identified 4 components that together explained 75% of the variance (Figure 4). We applied varimax rotation to the components to maximize the feature loading of each feature onto a single component. To test how the PCA axis mapped onto anatomical axes, we took the (x,y,z) anatomical position of each neuron and correlated it with the score of each component. We performed this in standard stereotaxic space (Supplementary Figure 8) and also for a rotated set of anatomical axis (axes were rotated along the x, y z, axis in 16 equally spaced rotations from 0 to pi). The rotated axis that best correlated with the scores of a component (maximal R-squared for linear correlation) was identified as the optimal axes for explaining that components variance.

For the supervised classification, we partitioned the subthalamic nucleus into two segments along the optimal axis (median split) and used these as labels to train a linear SVM classifier. We trained on a randomly selected subset of neurons (n= 9 neurons per region to give 27 total neurons; leaving out 1 neuron each from dorsal, 6 from central, and 3 from ventral) and tested on left-out data. We performed this procedure 1,000 times on different random subsets to obtain the average classification accuracy (chance=33%). To assess which features contributed the most to the classification we report the beta-coefficients of the SVM classifier for each binary comparison (e.g. ventral vs. central).

## Acknowledgements

We gratefully acknowledge the willingness of our patients to participate in this study. We thank Dr. Michele Tagliati for patient referrals, the staff and physicians of the Movement Disorders Program at Cedars-Sinai Medical Center for support; Robert Zelaya, Lori Scheinost, Andy Nguyen, and Cody Holland for assistance with intraoperative recordings; Jan Kaminksi, Jeong Woo Choi, and Steven Errington for discussion and feedback on the manuscript; Adam Aron for advice and for contributing to writing the grant that partially funded this work. This work was supported by the BRAIN initiative through grants from the National Institute of Neurological Disorders and Stroke (U01NS103792 to UR, U01NS098961 to NP and UR).

