## Supplementary Materials for "“Distinct roles of dorsal and ventral subthalamic neurons in action selection and cancellation.”"

**Table S1: Patient Demographics. (Related to Experimental Methods, Subject Details.)** ID=Subject identifier, M=Male, F=Female. Age at time of recording. Each patient performed the stop signal task between 1-4 times while recording neural activity in either the Left or Right STN. UPDRS III Scores are from pre-operative assessments, when the patient is off medication for Parkinson's Disease. (Note subject 86 did not have a pre-operative UPDRS score available, his post-operative score when off DBS and off Parkinson's medication was 16)

| Session # | ID | Surgery Side | Sex | Age | Ethnicity | UPDRS III Score |
|---|---|---|---|---|---|---|
| 1 | 65 | Left | M | 76 | White Non-Hispanic | 15 |
| 2 | 67 | Right | M | 72 | White Non-Hispanic | 55 |
| 3 | 68 | Left | F | 59 | Other Hispanic | 60 |
| 4 | | Right | | | | |
| 5 | 70 | Right | F | 65 | White Non-Hispanic | 44 |
| 6 | | Left | | | | |
| 7 | 71 | Right | F | 72 | White Non-Hispanic | 35 |
| 8 | 74 | Left | M | 71 | Other Non-Hispanic | 29 |
| 9 | | Right | | | | |
| 10 | 78 | Left | M | 72 | White Non-Hispanic | 26 |
| 11 | | | | | | |
| 12 | | Right | | | | |
| 13 | | | | | | |
| 14 | 80 | Left | F | 64 | White Non-Hispanic | 32 |
| 15 | 82 | Right | M | 69 | White Hispanic | 82 |
| 16 | 83 | Right | F | 47 | Asian Non-Hispanic | 13 |
| 17 | 85 | Right | M | 73 | White Non-Hispanic | 41 |
| 18 | | Left | | | | |
| 19 | 86 | Left | M | 79 | Other Non-Hispanic | 16* |
| 20 | | Right | | | | |
| 21 | 94 | Left | M | 75 | White Non-Hispanic | 21 |
| 22 | | Right | | | | |
| 23 | 96 | Left | F | 74 | White Non-Hispanic | 31 |
| 24 | | Right | | | | |
| 25 | 97 | Left | M | 59 | White Hispanic | 22 |
| 26 | | Right | | | | |
| 27 | | | | | | |
| 28 | 99 | Left | M | 72 | White Non-Hispanic | 33 |
| 29 | | | | | | |
| 30 | 100 | Right | M | 70 | White Non-Hispanic | 10* |
| 31 | 101 | Right | F | 62 | White Non-Hispanic | 38 |
| 32 | | | | | | |
| 33 | 102 | Left | F | 66 | White Hispanic | 31 |

* Pre-operative UPDRS scores off medication are not available. These are post-operative scores off medication and off DBS.

**Supplementary Figure 1 (Related to Experimental Methods, Spike Sorting): Spike Quality Sorting Metrics. (Related to Experimental Methods).** A-C. Histograms showing quality metrics of the 83 neurons: signal-to-noise ratio, spike amplitude, and percent of ISIs less than 3 ms for each neuron. Red line denotes mean. D. The % change in spike amplitude during stable trials for each neuron (on x-axis 0=first trial, 100% is last stable trial). E. The % change in firing rate during the stable trials.

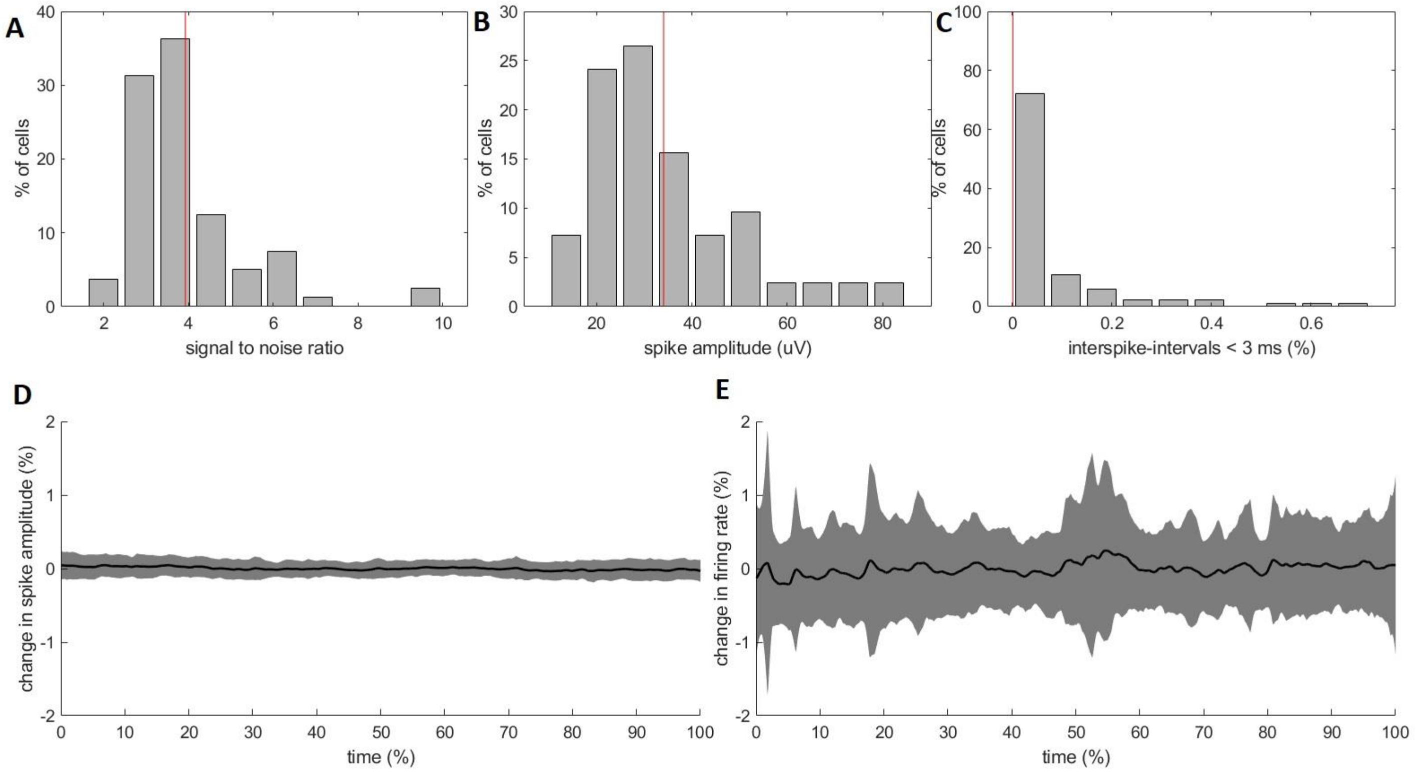

**Supplementary Figure 2 (Related to Figure 2): Example of other stop signal neurons.** Each panel shows the raster plot and PSTH for an individual stop signal neuron, activity aligned to the onset of the stop signal (or the time the stop signal would have occurred on a go-trial. Left panel compares successful stop trials (red) to latency matched slow go trials (dark blue). P-values show the statistical comparison (Wilcoxon rank sum test) of the binned firing rate (200 ms bin) at 1 ms intervals throughout the trial. Green lines denote time periods when the p-value is significant (<0.05) and survives cluster correction for multiple comparisons (see Methods). Right panels compare failed stop trials (orange) to latency matched fast go trials (light blue). A neuron was considered responsive to the stop-signal on successful or failed stop trials if the activity was significantly different from latency matched go-trials within 1 second of the presentation of the stop-signal.

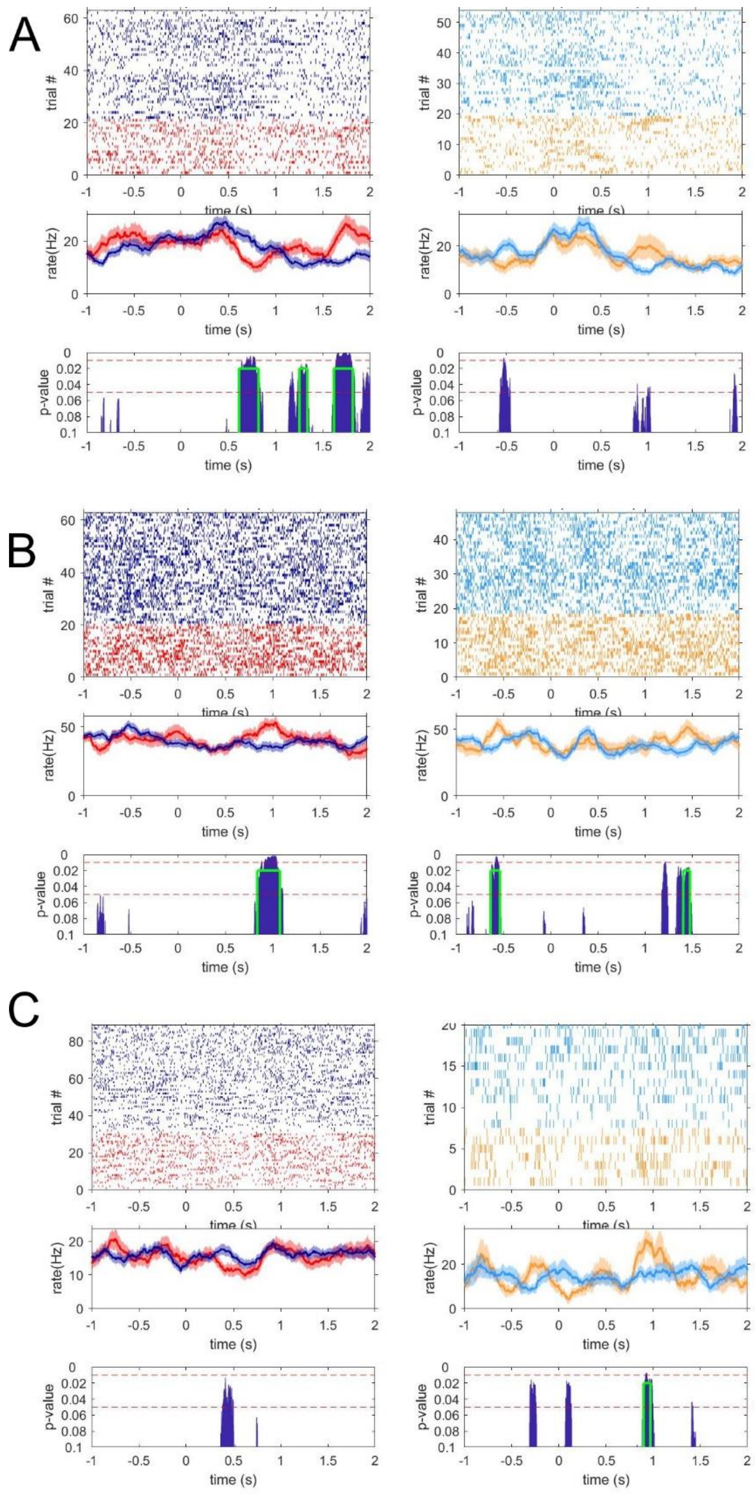

**Supplementary Figure 3: Spectrograms of LFP and ECog Power during the stop signal task (Related to Figure 3).** Spectrograms of the local field potential recorded from motor cortex (top) and STN (bottom). Left panels show the time-frequency spectrograms aligned to stop signal onset and averaged across all patients (z-scored relative to 500 ms pre-target baseline). Note that all trial types show beta suppression. Right panels show the difference in power between different trial types. Contour lines outline time-frequency points that are significantly different between conditions (paired t-test, p<0.05, cluster corrected for multiple comparisons). In both motor cortex and STN, there is less beta suppression during successful stopping; beta activity is elevated relative to slow-go trials.

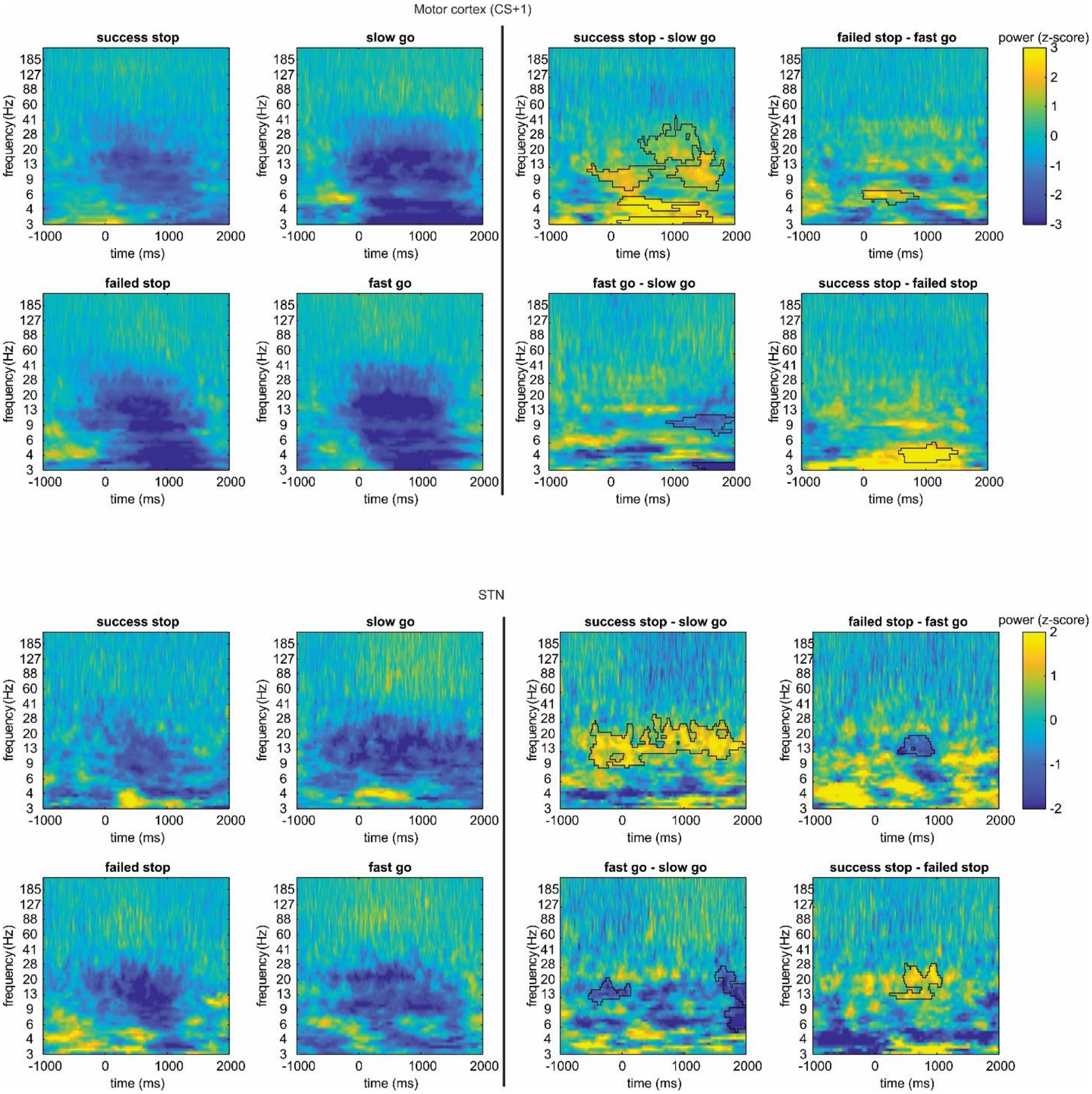

**Supplementary Figure 4 (Related to Figure 3): Spike-field associations between STN neurons and STN LFP and ECog iEEG.** (A) Example spike triggered average of the LFP signal in STN and the iEEG signal in motor cortex for a single neuron. The STA shows oscillatory activity in the beta band (13-30 Hz). (B) Percent of single neurons that significantly correlated with the STN LFP or ECog iEEG (significance assessed by comparing PPC, a measure of spike-field activity, to data where the LFP traces is randomly shifted in time, see Methods).

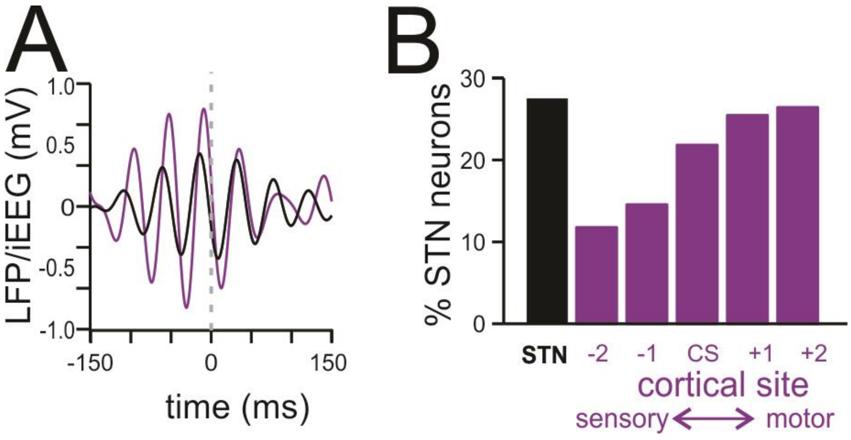

**Supplementary Figure 5 (Related to Figure 4): Spike Waveforms.** (A) Average spike waveforms of each STN neuron. Waveforms clustered into narrow (orange) and a broad (blue) groups based on the width of the waveform. (B) Distribution of waveform amplitude, half-width, trough-to-peak width, and repolarization time for the two groups (see methods). (C) The three width features of the two waveform groups. (D) The spike train features of each waveform group. Neurons with narrow and broad waveforms have significantly different activity during movement and in response to stop signals (see movement effect size and stop signal effect size, two sample t-test, $p<0.05$)

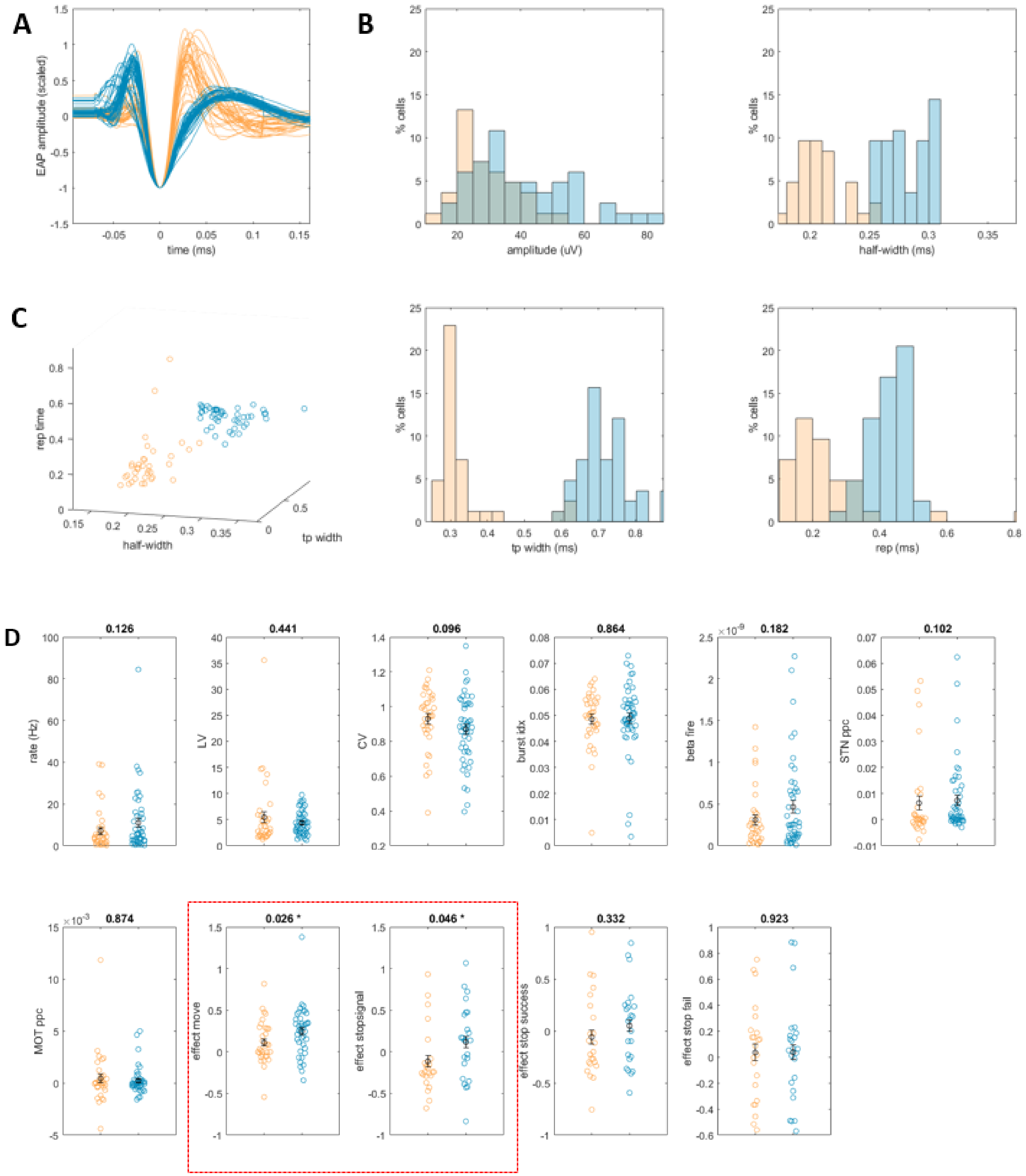

**Supplementary Figure 6 (Related to Figure 4): SVM classification using task features to decode dorsolateral vs ventromedial recording location along optimal anatomical axis.** Confusion matrix shows performance of a classifier used to decode dorsal vs. ventral anatomical location using only neural responses related to the task. Performance is significantly better than chance (72.8%, compared to 48% on shuffled labels, blue shaded region shows 95% confidence interval). Bar plot at bottom shows beta weights of the four firing rate features.

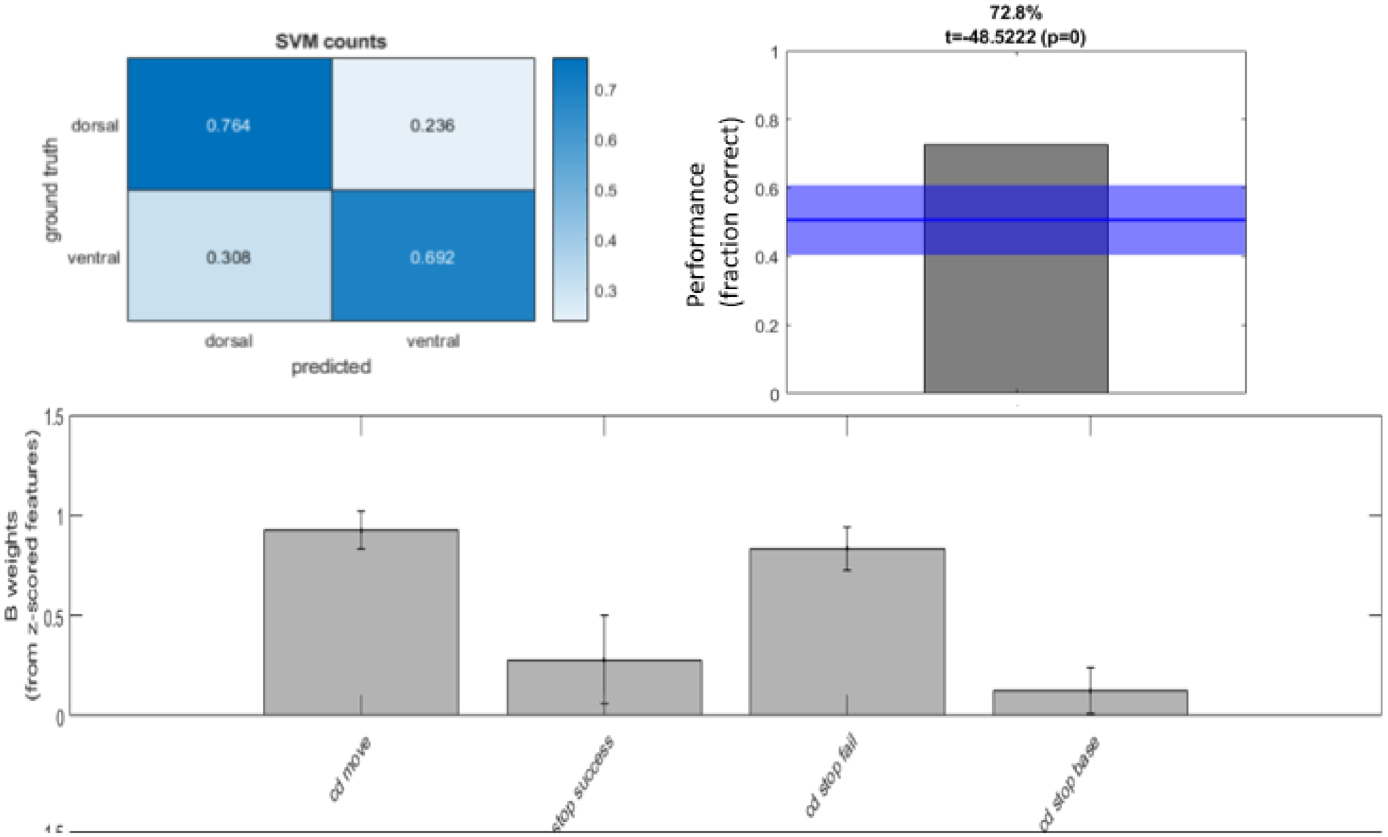

**Supplementary Figure 7 (Related to Figure 4): SVM classification of dorsal-ventral recording (MNI defined dorsal ventral axis).** Confusion matrix, performance, and beta weights plotted as in Supplementary Figure 6.

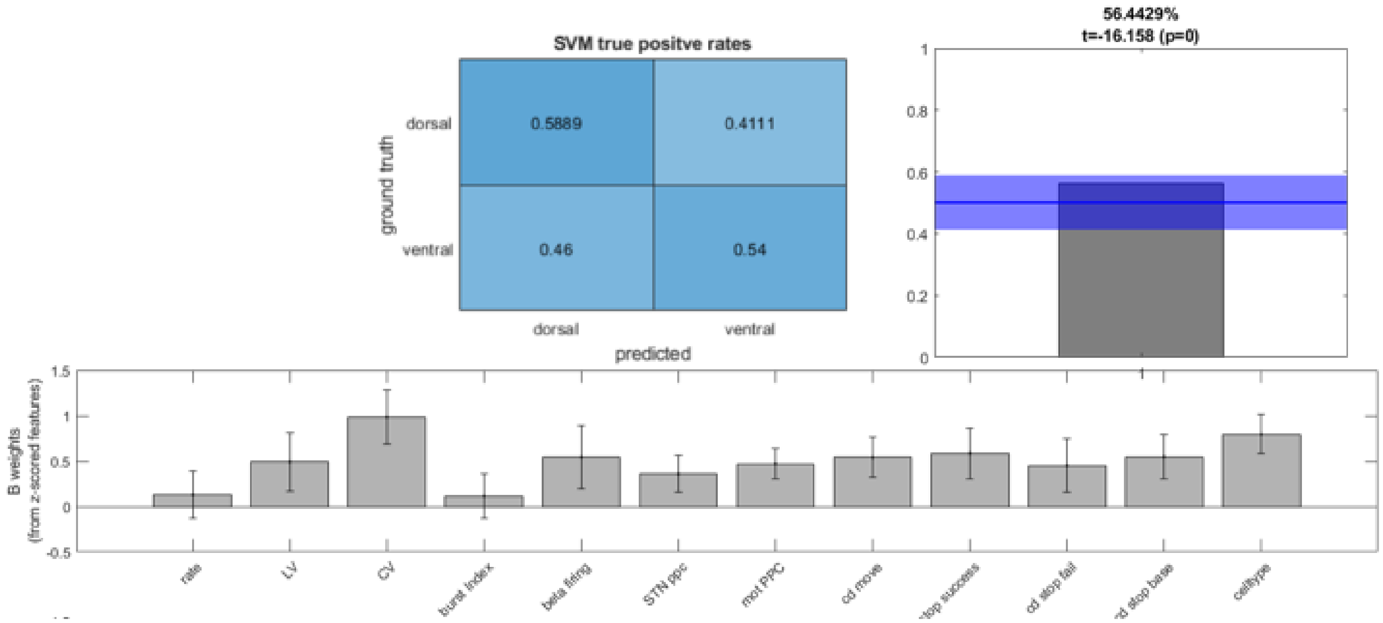

**Supplementary Figure 9 (Related to Experimental Methods, Subject Details) Distribution of firing features for patients with low vs. high UPDRS scores.** Patients were split into those with low vs. high UPDRS scores (median split). We calculated the average firing features across all neurons recorded from a single patient. Each dot represents a patient (green=low UPDRS score, pink =high). None of the features significantly differed between the two groups.

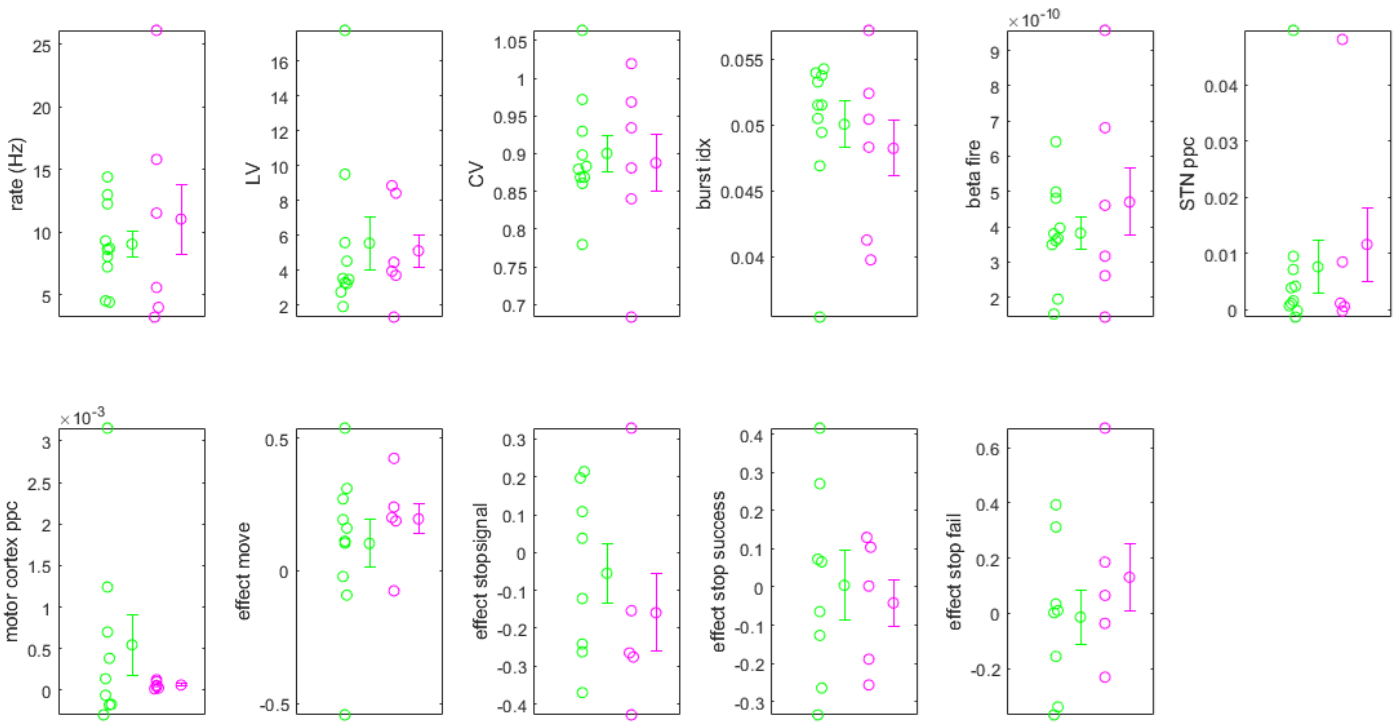
